# NPAS2 attenuates VSMC phenotypic switching in ascending thoracic aortic aneurysm via LPCAT3/PC-PUFA2S-mediated ferroptosis

**DOI:** 10.1101/2025.09.23.677952

**Authors:** Wenfeng Lin, Jiaqi Xiong, Yuhan Yan, Jinhui Bian, Zhen Zhang, Xinyang Xu, Yifei Diao, Xiaoxia Lan, Yu Zhong, Haibin Ji, Yongfeng Shao, Buqing Ni

## Abstract

Ascending thoracic aortic aneurysm (ATAA) is a life-threatening disorder with limited therapeutic options, critically linked to vascular smooth muscle cell (VSMC) phenotypic switching. Neuronal PAS Domain Protein 2 (NPAS2), a circadian rhythm-related transcription factor, regulates diverse physiological processes. However, the biological functions and underlying regulatory mechanisms of NPAS2 in VSMCs during ATAA pathogenesis remain to be elucidated. NPAS2 expression decreased in ATAA patients, PDGF-BB-treated HASMCs and BAPN-administrated mice. NPAS2 depletion facilitated VSMC phenotype switching. VSMC-specific NPAS2 knockout (NPAS2SMKO) mice exhibited aggravated ATAA. Mechanistically, NPAS2 depletion attenuates its transcriptional repression of LPCAT3, subsequently elevating the accumulation of phosphatidylcholines with two polyunsaturated fatty acyl chains (PC-PUFA2S), thereby promoting ferroptosis-induced VSMC phenotype switching and accelerating ATAA progression. Targeting NPAS2 represents a potential therapeutic strategy for ATAA treatment.

## Introduction

Ascending thoracic aortic aneurysm (ATAA) is a chronic aortic disease characterized by a localized dilation in the proximal segment of the aorta through maladaptive remodeling of the aortic wall. Without timely intervention, ATAA may progress to catastrophic complications, including aortic dissection or rupture, with a 24-hour mortality rate approaching 50%. ^1^Currently, pharmacological therapies exhibit limited efficacy in mitigating ATAA risk, while surgical intervention remains the optimal treatment option. ^2^ Hence, elucidating the regulatory mechanisms and developing novel therapeutic targets for ATAA remain urgent priorities.

Vascular smooth muscle cells (VSMCs), constituting the predominant cell types in the aorta, are essential for maintaining vascular tone homeostasis via vasoconstrictive and vasodilatory responses.^3, 4^ The phenotypic switching of VSMCs represents an adaptive response to various stress stimuli, including inflammation, oxidative stress, and hemodynamic alterations due to cardiovascular disease. Sustained stress stimuli induce synthetic transdifferentiation of contractile VSMCs, secreting elastolytic enzymes and proinflammatory cytokines to initiate pathological remodeling, thereby accelerating the ATAA progression.^1, 4^ Given its major pathogenic role, targeting molecular regulators of VSMC phenotypic transformation with cellular specificity is considered a promising therapeutic strategy.

The circadian clock serves as a vital endogenous regulator of physiological function. Oscillations of core clock genes critically sustain cardiovascular homeostasis.^5^ Neuronal PAS domain protein 2 (NPAS2), a heme-based sensor and DNA binding transcription factor,^6^ functions as an obligate heterodimer with BMAL1, that responds to diverse intra- and extracellular stimuli, playing a pivotal role in circadian rhythm regulation.^7^ In mammals, NPAS2 synchronizes metabolic processes with the body’s internal clock according to the day-night cycles. NPAS2 dysregulation disrupts circadian rhythms, thereby predisposing to sleep disturbances, neuropsychiatric conditions, and multisystem disorders including cerebro-cardiovascular diseases.^8^ For instance, NPAS2 correlates with the circadian phenotypes of hypertension, suggesting genetic modulation of blood pressure fluctuations in essential hypertension.^8^ However, the role of NPAS2 in ATAA remains uncharacterized.

The study elucidates the molecular mechanism by which NPAS2 depletion alleviates transcriptional repression of LPCAT3 to accumulate phosphatidylcholines with two polyunsaturated fatty acyl chains (PC-PUFA_2S_), thereby promoting ferroptosis-induced VSMC phenotype switching and accelerating ATAA progression.

Collectively, we identified NPAS2 as the potential therapeutic target for ferroptosis suppression, thereby proposing future research direction for ATAA treatment strategies.

## Methods

### Animal ethics statement

This study involving mice was approved by the Nanjing Medical University Animal Care and Use Committee (IACUC-2504030). All animal care and experimental protocols were approved by the Animal Research: Reporting of In Vivo Experiments (ARRIVE) guidelines.

### Cell culture

HASMCs were isolated from tissues of control donors (control-HASMCs) and ATAA donors (ATAA-HASMCs). HASMCs between passages 4 and 8 (P4-P8) were used in experiments. Briefly, the tunica media layer of the aorta was dissected into 1–2 mm3 sections and placed in T25 culture flasks. After tissue adhesion, sections were gently covered with 15 mL DMEM (15% FBS) and cultured in a 37°C incubator with 5% CO2. Explants were maintained for 5 days (unless the medium turned yellow earlier), with three-quarters of the DMEM replaced every 3 days. Cells migrated from the explants after 1-2 weeks. Explants were removed from the flask surface, and the adherent cells were harvested and expanded to passage 2 (P2).

### ATAA models

C57BL/6J mice were purchased from Jiangsu Laboratory Animal Center (Jiangsu, China.). C57BL/6J (male, 3 weeks old) mice were maintained on a standard diet and administered 0.5% β-aminopropionitrile (BAPN, Sigma-Aldrich, USA) dissolved in sterile saline (1 g/kg/day) for 4 weeks. Following the treatment period, mice were euthanized in a CO2 chamber followed by cervical dislocation. Aortic tissues were harvested and fixed in 4% paraformaldehyde for subsequent analysis.

### NPAS2^SMKO^ models

Tagln^Cre/+^ and NPAS2^fl/fl^ mice and were purchased from GemPharmatech Co. Ltd. Tagln^Cre/+^ mice were crossed with NPAS2^fl/fl^ mice to generate VSMC-specific ablation of NPAS2 (NPAS2^SMKO^). PCR primers (5′–3′) used for genotyping were as follows: TGCCACGACCAAGTGACAGCAATG and ACCAGAGACGGAAATCCATCGCT C (Tagln^Cre/+^ mice); 5’arm CAAATAGGACAGCATCAGTGTCTGTC/CCGCCTCT CTAACTTTGTAGAGCTTT and 3’arm CATCGCATTGTCTGAGTAGGTG/GTCA TGGTCACATGGCAGGATAA (NPAS2^fl/fl^ mice). All mice were on a C57BL/6J background, maintained under specific pathogen-free conditions, with free access to a standard mouse chow and sterile water unless otherwise specified.

### Immunofluorescence

HASMCs were fixed with prechilled absolute ethanol for 10-20 min at 60–70% confluency. After three PBS washes, HASMCs were permeabilized and blocked in 0.3% Triton X-100 containing 5% bovine serum albumin (BSA) for 1 h at 25°C, followed by three additional PBS washes. Primary antibodies diluted in blocking buffer were then applied and incubated overnight at 4°C, followed by three PBS washes. HASMCs were incubated with fluorophore-conjugated secondary antibodies in the dark at 25°C for 40 min. After three PBS washes, HASMCs were mounted with DAPI to stain nuclei. Images were acquired by a Thunder DMi8 Fluorescence Microscope (Leica Microsystems, Germany). The antibodies used are listed in Supplemental Table 1.

### Chromatin Immunoprecipitation

Chromatin immunoprecipitation (ChIP) was performed using the EZ-ChIP kit (17-10086; Millipore). Briefly, cells were cross-linked with 1% formaldehyde for 10 minutes and harvested in SDS lysis buffer, followed by sonication to shear chromatin. Cross-linked protein/DNA was immunoprecipitated with anti-NPAS2 (365666X; Santa Cruz Biotechnology), anti-GFP (A-11120; Invitrogen), or control immunoglobulin G. Promoter binding was determined by quantitative PCR.

### Quantitative real-time PCR

Total RNA was extracted from aortic tissues and VSMCs using TRIzol^TM^ Reagent (Thermo Fisher Scientific, USA) according to the manufacturer’s instructions. Reverse transcription and cDNA synthesis were performed using the HiScript lll RT SuperMix for qPCR (Vazyme Biotech, China). Quantitative Real-time PCR was performed with SYBR qPCR Master Mix (Vazyme), and data were collected using QuantStudio Real-Time PCR Software (Thermo). Relative gene expression was calculated using the 2ΔΔCt method with GAPDH as a reference. Primer sequences used are listed in Supplemental Table 2.

### Western blot analysis

Tissues and VSMCs were lysed with RIPA buffer containing protease and phosphatase inhibitors (NCM Biotech, China), and incubated on ice for 30 min. Lysates were centrifuged at 13,800g for 30 min at 4°C, and supernatants were collected. Protein concentration was quantified using a bicinchoninic acid protein assay kit (Beyotime, China) according to the manufacturer’s instructions. The proteins were denatured in 5×SDS loading buffer (Sangon Biotech, China) at 100 °C for 8 min. The experimental and control samples co-electrophoresed on SDS-PAGE gels and electroblotted onto PVDF membranes (Roche, Switzerland) using the wet-transfer system. Membranes were activated in methanol prior to transfer. After blocking with 6% (wt/vol) skim milk in TBST for 2 h at 26°C, membranes were sectioned according to molecular weight markers and incubated with primary antibodies overnight at 4°C. Following five 5-min TBST washes, membranes were incubated with corresponding horseradish peroxidase (HRP)-conjugated secondary antibodies (ZSGB-BIO, China) for 1.5 h at 26°C. After additional washes, protein bands were visualized using ECL reagent (Thermo). When required, membranes were stripped with Restore™ PLUS Western Blot Stripping Buffer (Thermo) and re-probed. Protein levels were normalized to GAPDH levels. Protein quantification was performed using ImageJ software (NIH, USA). The antibodies are listed in Supplementary Table 3.

### Gel zymography

Supernatants harvested from cultured HASMCs underwent centrifugation to generate concentrated conditioned media, as previously described^4^. Protein concentration was normalized using a BCA assay. The conditioned media were electrophoresed on 10% SDS-PAGE co-polymerized with 1 mg/mL gelatin (as a substrate for MMP activity). Gels were washed 2 times with 2.5% (v/v) Triton X-100 to renature gelatinases, and then equilibrated in developing buffer (50 mM Tris-HCl [pH 7.4], 150 mM NaCl, 5 mM CaCl2, and 0.1% Brij-35) for 48 h at 37°C. Gels were then stained using 0.25% Coomassie brilliant blue for 2 h and destained in 30% methanol/10% acetic acid until clear bands appeared against a blue background. Gelatinolytic activity was quantified using ImageJ software (NIH).

### Collagen gel contraction assay

HASMCs were cultured in serum-free medium for 24 h, then embedded into collagen gels, as previously described^4^. Briefly, collagen gels were prepared by mixing HASMCs with type I collagen (Thermo), 2×DMEM, sterile 1 M NaOH, and distilled water. The collagen-cell suspension was immediately dispensed in cell culture plates and incubated for 30 min at 37°C. Following polymerization, gels were then gently released from the well surfaces using pipette tips. Contraction kinetics were documented by capturing digital images, and the gel surface areas were quantified using ImageJ.

### Human ethics statement, patient selection and specimen acquisition

This study involving human participants was approved by the Ethics Committee of the First Affiliated Hospital with Nanjing Medical University (IRB number:2019-SR-067). All procedures complied with the ethical standards stated in the Declaration of Helsinki. Written informed consent was obtained from all participants preoperatively. Experimental specimens (N =6) were collected from ascending thoracic aortic aneurysm patients undergoing aortic aneurysm repair at our institution. Samples were taken from the dilated ascending aneurysms of patients. Control specimens (N = 6) were obtained from age-matched patients undergoing heart transplantation.

### Statistical analysis

Data analysis was performed using GraphPad Prism 9.4.1 (GraphPad Software, La Jolla, CA, USA). Data are expressed as mean ± standard deviation. Differences were analyzed by the unpaired t-test, one-way analysis of variance or Kaplan-Meier. P < 0.05 was deemed to indicate statistical significance.

## Results

### Decreased NPAS2 is associated with ATAA tissues

To investigate potential genes associated with VSMC phenotypic switching in ascending thoracic aortic aneurysm (ATAA), we initially analyzed the gene expression profiles of human ATAA (GSE26155), mouse ATAA (GSE241968) from Gene Expression Omnibus database to identify ATAA-related transcriptomic hallmarks. We next sought to identify circadian rhythm-related genes in mouse ascending thoracic aortas (GSE54650, GSE54651). Venn diagram analysis illustrated global overlap of differentially expressed genes, with NPAS2 and VGLL3 emerging as statistically significant circadian rhythm-associated genes in ATAA (Figure 1A). To validate this finding, we analyzed 12 human specimens, including 6 ATAA tissues and 6 control tissues. qPCR analysis showed significantly reduced NPAS2 mRNA expression in ATAA specimens compared to control specimens (Figure 1B). Western blot experiments showed that NPAS2 expression was decreased in ATAA tissues compared with normal tissues, along with increased levels of the synthetic proteins (OPN and KLF4) and decreased levels of contractile proteins (α-SMA and SM22-α) (Figure 1C). Immunofluorescence showed decreased NPAS2 and α-SMA expression in ATAA tissues versus normal tissues (Figure 1D). These results suggest NPAS2 plays a crucial role in ATAA. Since VSMC phenotype switching is a major contributor to ATAA pathogenesis, we investigated whether NPAS2 participates in VSMC phenotype switching. Following PDGF-BB treatment, Western blot analysis demonstrated reduced NPAS2 expression accompanied by elevated synthetic proteins (OPN, KLF4) and diminished contractile proteins (α-SMA, SM22-α) (Figure 1E). Immunofluorescence analysis demonstrated reduced NPAS2 and α-SMA expression after PDGF-BB treatment (Figure 1G). Conversely, OPN and KLF4 protein levels decreased, whereas NPAS2, α-SMA and SM22-α protein levels increased after TGF-β1 treatment (Figure 1F). Immunofluorescence staining showed enhanced NPAS2 and α-SMA expression following TGF-β1 treatment (Figure 1G).

**Figure 1.**
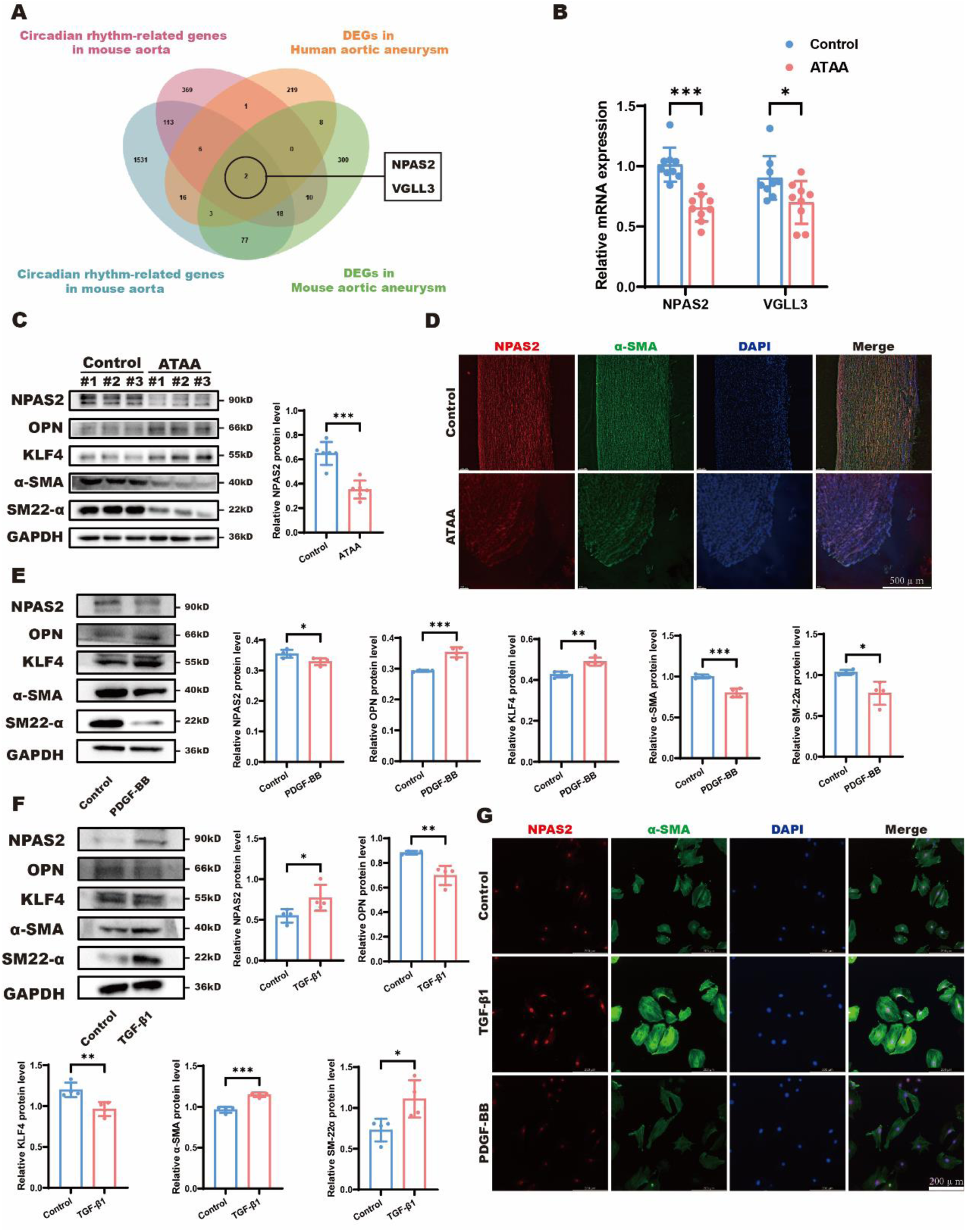
NPAS2 negatively correlates with VSMC phenotypic switching in ATAA tissues. (A) Venn diagram depicting the intersection of circadian rhythm-related genes in mouse aorta (GSE54650, GSE54651), differentially expressed genes (DEGs) in human ATAA (GSE26155), and DEGs in mouse ATAA (GSE241968). (B) Quantification of NPAS2 and VGLL3 mRNA levels in control and ATAA tissues. n = 6. (C) Representative immunoblots and quantification of NPAS2, OPN, KLF4, α-SMA, and SM22-α in control and ATAA tissues. n = 6. (D) Representative immunofluorescence images of NPAS2 (red), α-SMA (green), and DAPI (blue) in control and ATAA tissues. Scale bars: 500 μm. n = 6. (E) Representative immunoblots and quantification of NPAS2, OPN, KLF4, α-SMA, and SM22-α in control and PDGF-BB-treated HASMCs. n = 4. (F) Representative immunoblots and quantification of NPAS2, OPN, KLF4, α-SMA, and SM22-α in control and TGF-β1-treated HASMCs. n = 4. (G) Representative immunofluorescence images of NPAS2 (red), α-SMA (green), and DAPI (blue) in control, TGF-β1-treated and PDGF-BB-treated HASMCs. Scale bars: 200 μm. n = 6. *p < 0.05, **p< 0.01, ***p< 0.001. B, E and F, unpaired two-tailed t-test; C, one-way ANOVA.

### NPAS2 inhibits phenotypic switching of VSMC to delay ATAA

Given our finding that NPAS2 negatively correlates with VSMC phenotypic transformation, we further explored whether NPAS2 inhibits VSMC phenotypic switching to delay ATAA progression. Following transfection of HASMCs with NPAS2 siRNA (si-NPAS2), OPN and KLF4 levels increased, whereas α-SMA and SM22-α levels decreased (Figure 2A). Immunofluorescence staining showed reduced α-SMA expression in si-NPAS2-treated HASMCs (Figure 2B). Gelatin zymography assays depicted enhanced MMP-9 and MMP-2 activities in si-NPAS2-transfected HASMCs (Figure 2C). Cell contraction assays demonstrated NPAS2 knockdown impaired HASMC contractility (Figure 2D). Conversely, OPN and KLF4 levels decreased while α-SMA and SM22-α levels increased, respectively, after HASMCs were transfected with NPAS2 over-expression plasmid (oe-NPAS2) (Figure 2E). Increased α-SMA expression in oe-NPAS2-treated HASMCs was observed via immunofluorescence (Figure 2F). Decreased MMP-9 and MMP-2 activities in oe-NPAS2-transfected HASMCs were demonstrated by gelatin zymography (Figure 2G). NPAS2 overexpression enhanced HASMC contractility as measured by cell contraction assays (Figure 2H).

**Figure 2.**
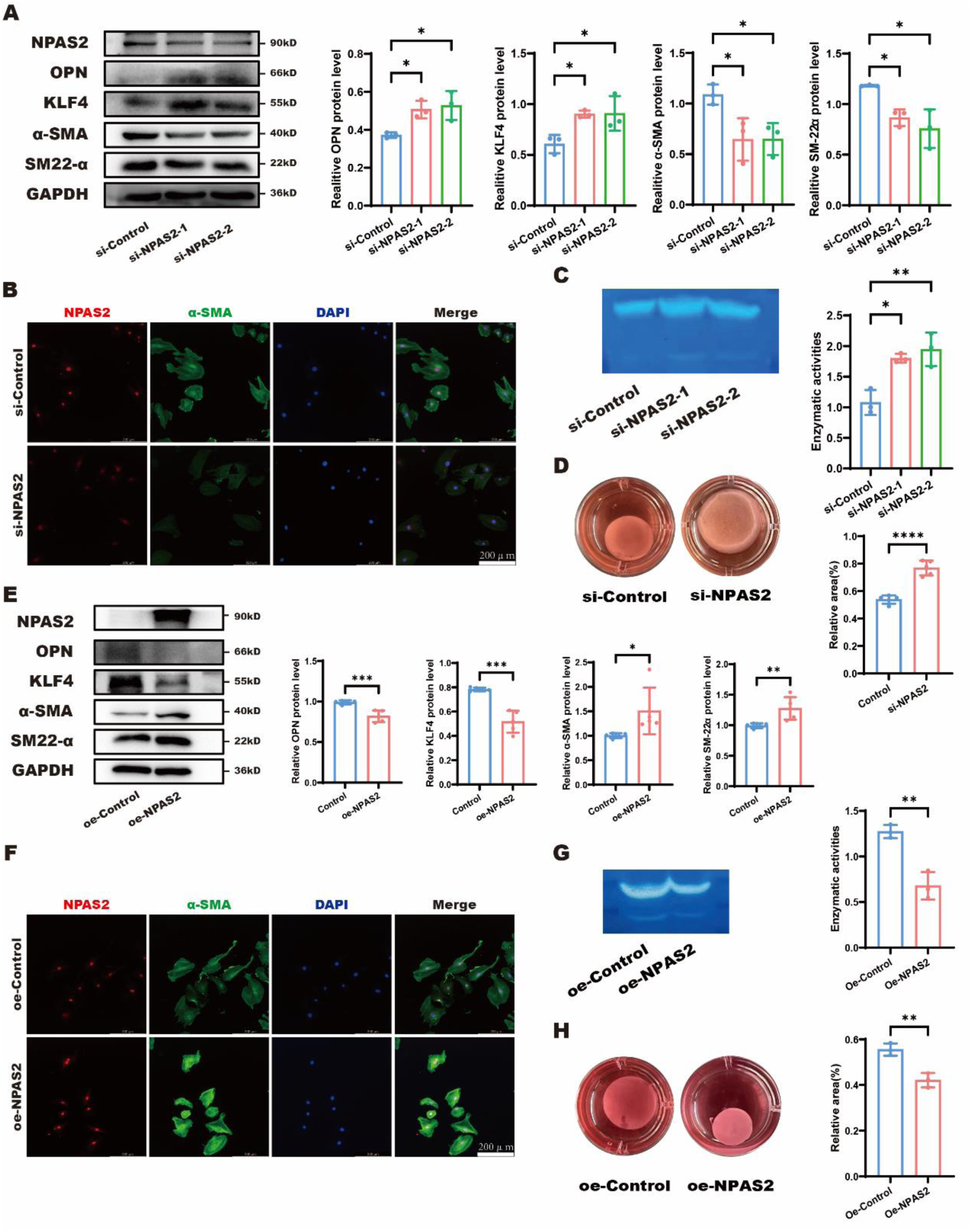
NPAS2 inhibits phenotypic switching of HASMCs. (A) Representative immunoblots and quantification of NPAS2, OPN, KLF4, α-SMA, and SM22-α in HASMCs transfected with control siRNA (si-Control) or NPAS2-targeting siRNA (si-NPAS2). n =3. (B) Representative immunofluorescence images of NPAS2 (red), α-SMA (green), and DAPI (blue) in si-Control- or si-NPAS2-transfected HASMCs. Scale bars: 200 μm. n = 6. (C) Representative gelatin zymography images and quantification of MMP-2 and MMP-9 enzyme activity in si-Control- or si-NPAS2-transfected HASMCs. n = 5. (D) Representative contraction images and quantification of Collagen I in si-Control- or si-NPAS2-transfected HASMCs. Scale bars: 5 mm. n = 5. (E) Representative immunoblots and quantification of NPAS2, OPN, KLF4, α-SMA, and SM22-α in HASMCs transfected with empty plasmid (oe-Control) or NPAS2 over-expression plasmid (oe-NPAS2). n = 5. (F) Representative immunofluorescence images of NPAS2 (red), α-SMA (green), and DAPI (blue) in oe-Control- or oe-NPAS2-transfected HASMCs. Scale bars: 200 μm. n = 6. (G) Representative gelatin zymography images and quantification of MMP-2 and MMP-9 enzyme activities in oe- Control- or oe-NPAS2-transfected HASMCs. n = 5. (H) Representative contraction images and quantification of Collagen I in oe-Control- or oe-NPAS2-transfected HASMCs. Scale bars: 5 mm. n = 5. *p < 0.05, **p< 0.01, ***p< 0.001, ****p< 0.0001. A and C, one-way ANOVA; D, E, G and H, unpaired two-tailed t-test.

To confirm NPAS2 negatively correlates with ATAA in vivo, mice with specific NPAS2 depletion in VSMCs were generated (NPAS2^SMKO^). An ascending thoracic aortic aneurysm model was then established in 3-week-old male NPAS2^WT^ and NPAS2^SMKO^ mice through 4-week BAPN administration (Figure 3A). NPAS2^SMKO^ mice treated with BAPN had a lower survival probability than BAPN-treated NPAS2^WT^ mice (Figure 3B). As anticipated, BAPN-treated mice developed thoracic aortic pathology (dilation or aneurysm), as confirmed through gross and ultrasonographic examination. Compared with BAPN-treated NPAS2^WT^ mice, BAPN-treated NPAS2^SMKO^ mice exhibited significantly higher susceptibility to ATAA formation, as evidenced by larger thoracic aortic dilations/aneurysms on gross examination (Figure 3C), greater maximal internal aortic diameters on ultrasonography (Figure 3D), more severe disruption of vascular architecture in H & E staining, increased fragmentation and degradation of elastic fibers in EVG staining, worse degradation of muscle fibers and more hyperplasia of collagen fibers in Masson staining (Figure 3E), and decreased α-SMA expression in immunofluorescence (Figure 3F). These aforementioned results confirmed the potent inhibitory effect of NPAS2 on VSMC phenotype switching in ATAA progression.

**Figure 3.**
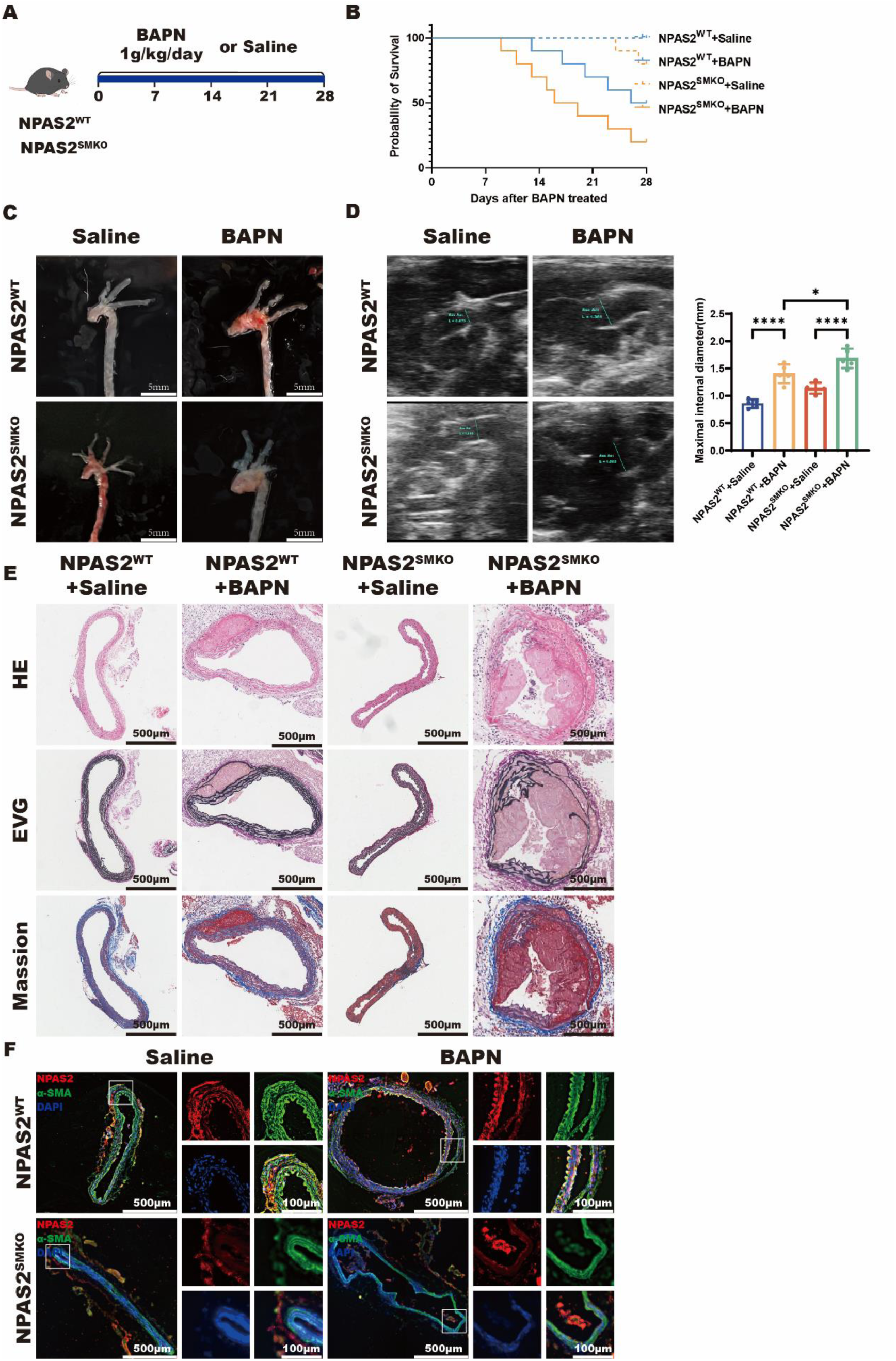
NPAS2 knockout in VSMCs aggravates ATAA in BAPN-treated mice. (A) NPAS2^WT^ mice and NPAS2^SMKO^ mice(male, 3 weeks old) were fed a standard diet and administered saline or 0.5% β-aminopropionitrile (BAPN) (1g/kg/day) for 28 days. n = 10. (B) Survival probability of NPAS2^WT^ mice and NPAS2^SMKO^ mice administered saline or BAPN at 7, 14, 21 and 28 days. n = 10. (C) Representative morphology of ascending thoracic aortas in NPAS2^WT^ mice and NPAS2^SMKO^ mice after BAPN administration. Scale bars: 5 mm. n = 10. (D) Representative ultrasound images and inner diameter quantification of ascending thoracic aortas in NPAS2^WT^ mice and NPAS2^SMKO^ mice after BAPN administration. n = 8-10. (E) Representative H&E, EVG and Masson staining of ascending thoracic aortas in NPAS2^WT^ mice and NPAS2^SMKO^ mice after BAPN administration. Scale bars: 500 μm. n = 6. (F) Representative immunofluorescence images of NPAS2 (red), α-SMA (green), and DAPI (blue) in NPAS2^WT^ mice and NPAS2^SMKO^ mice after BAPN administration. Scale bars:500 μm. n = 6. *p < 0.05, ****p< 0.0001. B, Kaplan-Meier; D, one-way ANOVA.

### NPAS2 deficiency-induced accumulation of PLs promotes VSMC phenotypic switching

Next, we sought to further investigate the physiological relevance of NPAS2-mediated phenotypic switching in HASMCs. Transcriptome RNA sequencing of HASMCs with or without NPAS2 knockdown revealed profound and statistically significant alterations in lipid metabolism (Figure 4A). HASMCs accumulated higher levels of neutral lipids after transfected with si-NPAS2, as evidenced by BODIPY^493/503^ staining (Figure 4B). To elucidate the mechanism underlying how NPAS2-mediated regulation of phenotypic switching, we sought to analyze alterations in lipid profiles caused by NPAS2 depletion. Lipidomic liquid chromatography–tandem mass spectrometry (LC-MS/MS) lipidomic analysis of si-Control- and si-NPAS2-transfected HASMCs revealed significantly increased total lipid content upon NPAS2 depletion (Figure 4C).We further screened out and identified the genes with statistically significant alterations, including phosphatidylethanolamine (PE), Ceramide (Cer), Monoglyceride (MG), Lysophosphatidylcholine (LPC), fatty acids (FA), Diacylglycerol (DG), phosphatidylcholine (PC) and Phosphoinositide (PI) (Figure 4 D, E).Next, we aimed to identify specific lipids among the aforementioned candidates that drive VSMC phenotypic transition. HASMCs were treated with exogenous lipids, including arachidonic acid, 1-Monoarachidin, LysoPC (20:4), PC (20:4,20:4), respectively. PC (20:4,20:4) exhibited the most pronounced specificity and efficacy in inducing phenotypic switching. This effect was evidenced by the highest OPN, KLF4 protein levels, the lowest α-SMA, SM22-α protein levels in immunoblot (Figure 4F), the strongest MMP-2 and MMP-9 activities in gelatin zymography (Figure 4G), the minimal contractility in collagen gel contraction assay (Figure 4H). Thus, phospholipids induce phenotypic switching in HASMCs.

**Figure 4.**
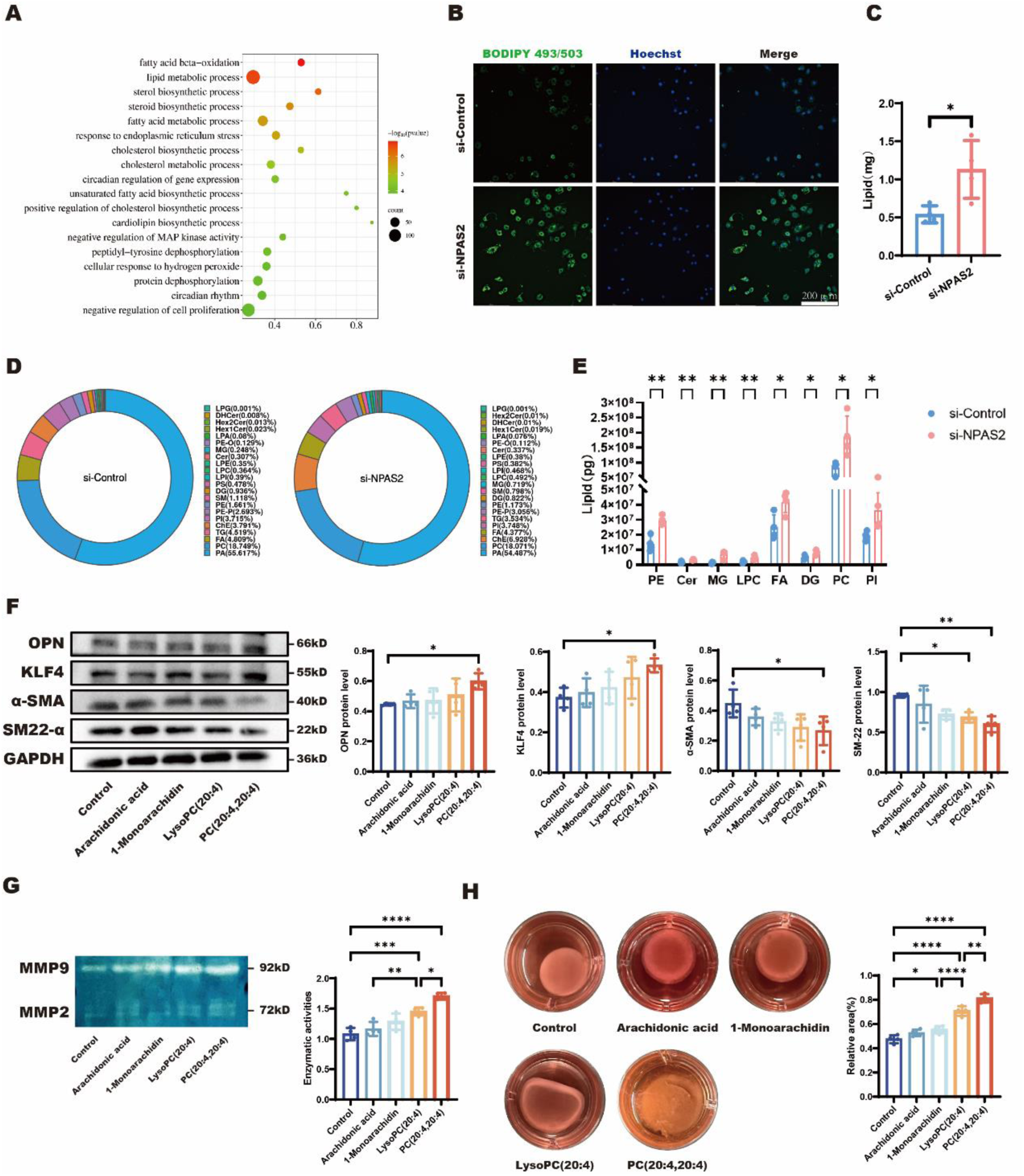
PLs induces phenotypic switching in HASMCs. (A) Transcriptome RNA sequencing analysis of HASMCs with and without NPAS2 knockdown. n = 4. (B) Representative immunofluorescence images of lipid droplets in si-Control and si-NPAS2 HASMCs. Scale bars: 200 μm. n = 6. (C) Lipidomic liquid chromatography–tandem mass spectrometry (LC-MS/MS) analysis of total lipid content in si-Control and si-NPAS2 HASMCs. n = 4. (D) LC-MS/MS analysis of statistically significant lipid species in si-Control and si-NPAS2 HASMCs. n = 4. (E) LC-MS/MS analysis identified eight significantly altered lipid species (PE, Cer, MG, LPC, FA, DG, PC and PI) in si-Control and si-NPAS2 HASMCs. n = 4. (F) Representative immunoblots and quantification of OPN, KLF4, α-SMA, and SM22-α in HASMCs treated with Arachidonic acid, 1-Monoarachidin, LysoPC (20:4), PC (20:4, 20:4), respectively. n = 4. (G) Representative gelatin zymography images and quantification of MMP-2 and MMP-9 enzyme activities in HASMCs treated with Arachidonic acid, 1-Monoarachidin, LysoPC (20:4), PC (20:4, 20:4), respectively. n = 5. (H) Representative contraction images and quantification of Collagen I in HASMCs treated with Arachidonic acid, 1-Monoarachidin, LysoPC (20:4), PC (20:4,20:4), respectively. Scale bars: 5 mm. n = 5. * p < 0.05, **p< 0.01, ****p< 0.0001. C and E, unpaired two-tailed t-test; F and G, one-way ANOVA.

To elucidate the key phospholipids driving VSMC phenotypic switching among statistically significant candidates, we treated HASMCs with exogenous lipids, including arachidonic acid, Diarachidonin (20:4,20:4), PC (20:4,20:4), PI (20:4,20:4), PE (20:4,20:4), Cer (20:4,20:4), respectively. PC (20:4,20:4) showed significantly greater selectivity and potency than other PLs in inducing phenotypic switching, as confirmed by higher synthetic markers (Figure 5A), higher MMP-2 and MMP-9 activities (Figure 5B), and less HASMC contractility (Figure 5C). PC, the most significant phospholipids, induce phenotypic switching in HASMCs. For PCs, the sn1 and sn2 positions of the glycerol backbone can contain different fatty acids with varying degrees of unsaturation, saturated fatty acyl (SFA), monounsaturated fatty acyl (MUFA), or polyunsaturated fatty acyl (PUFA). To determine whether fatty acid saturation influences phospholipid-mediated regulation of VSMC phenotype, we investigated the effect of NPAS2 depletion on fatty acid saturation profiles. NPAS2 downregulation reduced SFA content while increasing PUFA content in PC species (Figure 5D). Alterations in other phospholipid classes are presented in Supplemental Figure S1. To explore the functional impact of the polyunsaturated fatty acyl chain in PCs on VSMC phenotypic switching, we treated HASMCs with PC (18:0,18:0), PC (18:0,20:4), and PC (20:4,20:4). PCs with two polyunsaturated fatty acyl chains (PC-PUFA_2S_) more potently induced VSMC phenotypic switching than PCs with one polyunsaturated fatty acyl chain (PC-PUFA_1S_), as evidenced by higher OPN, KLF4 protein levels, lower α-SMA, SM22-α protein levels (Figure 5E), stronger MMP-2 and MMP-9 activities (Figure 5F), and less HASMC contractility (Figure 5G).

**Figure 5.**
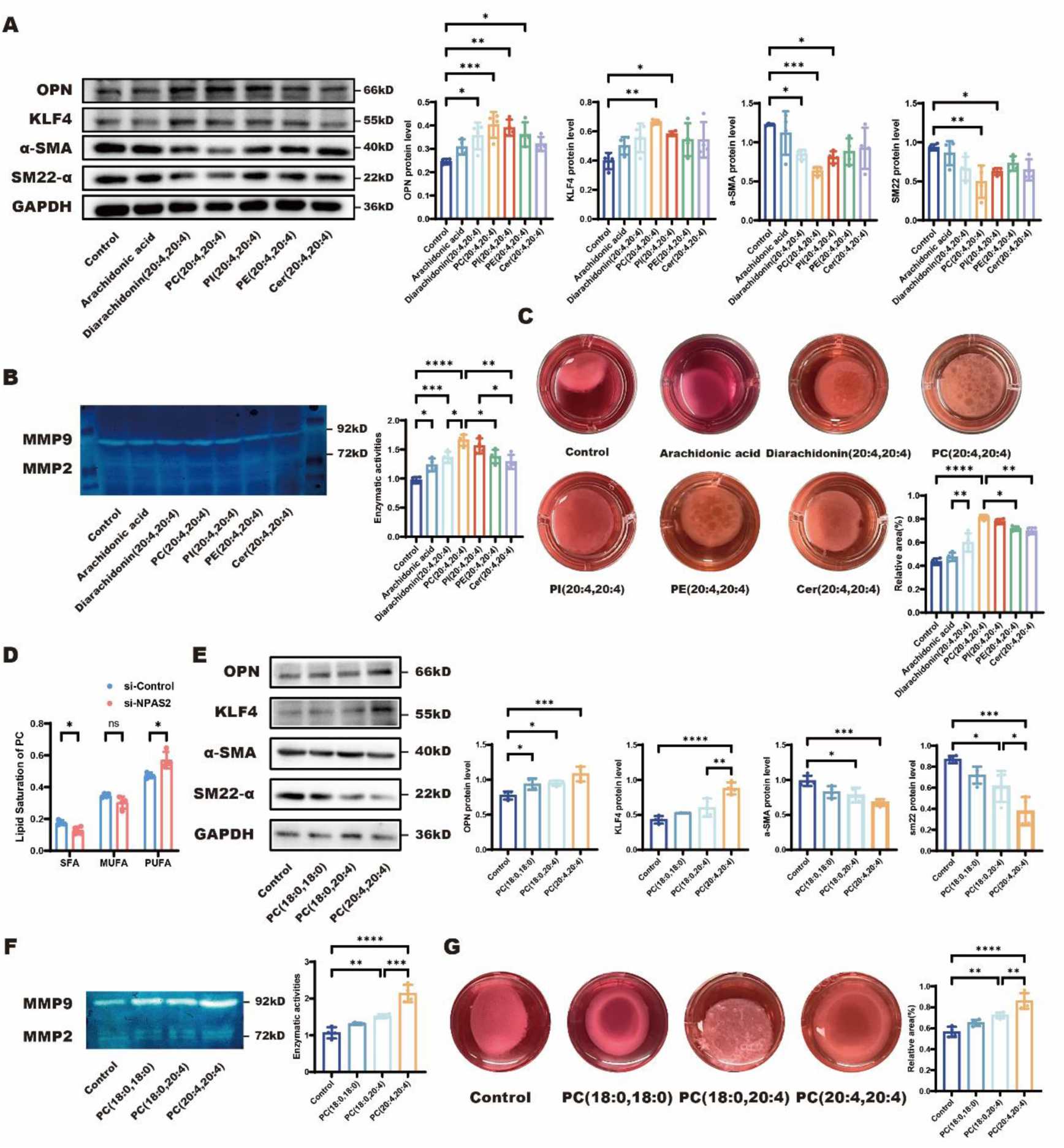
PC with two PUFA chains (PC-PUFA_2S_) exhibited more pronounced potency in inducing phenotypic switching of HASMCs. (A) Representative immunoblots and quantification of OPN, KLF4, α-SMA, and SM22-α in HASMCs treated with Arachidonic acid, Diarachidonin (20:4, 20:4), PC (20:4, 20:4), PI (20:4, 20:4), PE (20:4, 20:4), Cer (20:4, 20:4), respectively. n = 4. (B) Representative gelatin zymography images and quantification of MMP-2 and MMP-9 enzyme activities in HASMCs treated with Arachidonic acid, Diarachidonin (20:4, 20:4), PC (20:4, 20:4), PI (20:4, 20:4), PE (20:4, 20:4), Cer (20:4, 20:4), respectively. n = 5. (C) Representative contraction images of Collagen I in HASMCs treated with Arachidonic acid, Diarachidonin (20:4, 20:4), PC (20:4, 20:4), PI (20:4, 20:4), PE (20:4, 20:4), Cer (20:4, 20:4), respectively. Scale bars: 5 mm. n = 5. (D) Representative immunoblots and quantification of OPN, KLF4, α-SMA, and SM22-α in HASMCs treated with PC (18:0, 18:0), PC (18:0, 20:4), PC (20:4, 20:4), respectively. n = 4. (E) Representative gelatin zymography images and quantification of MMP-2 and MMP-9 enzyme activities in HASMCs treated with PC (18:0, 18:0), PC (18:0, 20:4), PC (20:4, 20:4), respectively. n = 5. (F) Representative contraction images of Collagen I in HASMCs treated with PC (18:0, 18:0), PC (18:0, 20:4), PC (20:4, 20:4), respectively. Scale bars: 5 mm. n = 5. *p < 0.05, **p< 0.01, ***p< 0.001, ****p< 0.0001. A, B, C, E, F and G, one-way ANOVA; D, unpaired two-tailed t-test.

To confirm the mechanistic role of PUFA_2S_ in VSMC phenotypic switching in vivo, NPAS2^WT^ or NPAS2^SMKO^ mice were treated with saline or arachidonic acid (AA) during BAPN administration (Figure 6A). Mice administered both AA and BAPN administration showed poor survival probability compared to those administered BAPN alone (Figure 6B). Compared to mice treated with BAPN, mice treated with both AA and BAPN developed more severe ATAA, characterized by larger thoracic aortic dilations/aneurysms on gross examination (Figure 6C), greater maximal internal diameters of aortas on ultrasonography (Figure 6D), more severe disruption of vascular architecture in H & E staining, increased fragmentation and degradation of elastic fibers in EVG staining, worse muscle fiber degradation and increased collagen fiber hyperplasia in Masson staining (Figure 6E), and reduced α-SMA expression in immunofluorescence (Figure 6F). The aforementioned results indicate that PUFA_2S_ exacerbated BAPN-induced ATAA in both NPAS2^WT^ and NPAS2^SMKO^ mice.

**Figure 6.**
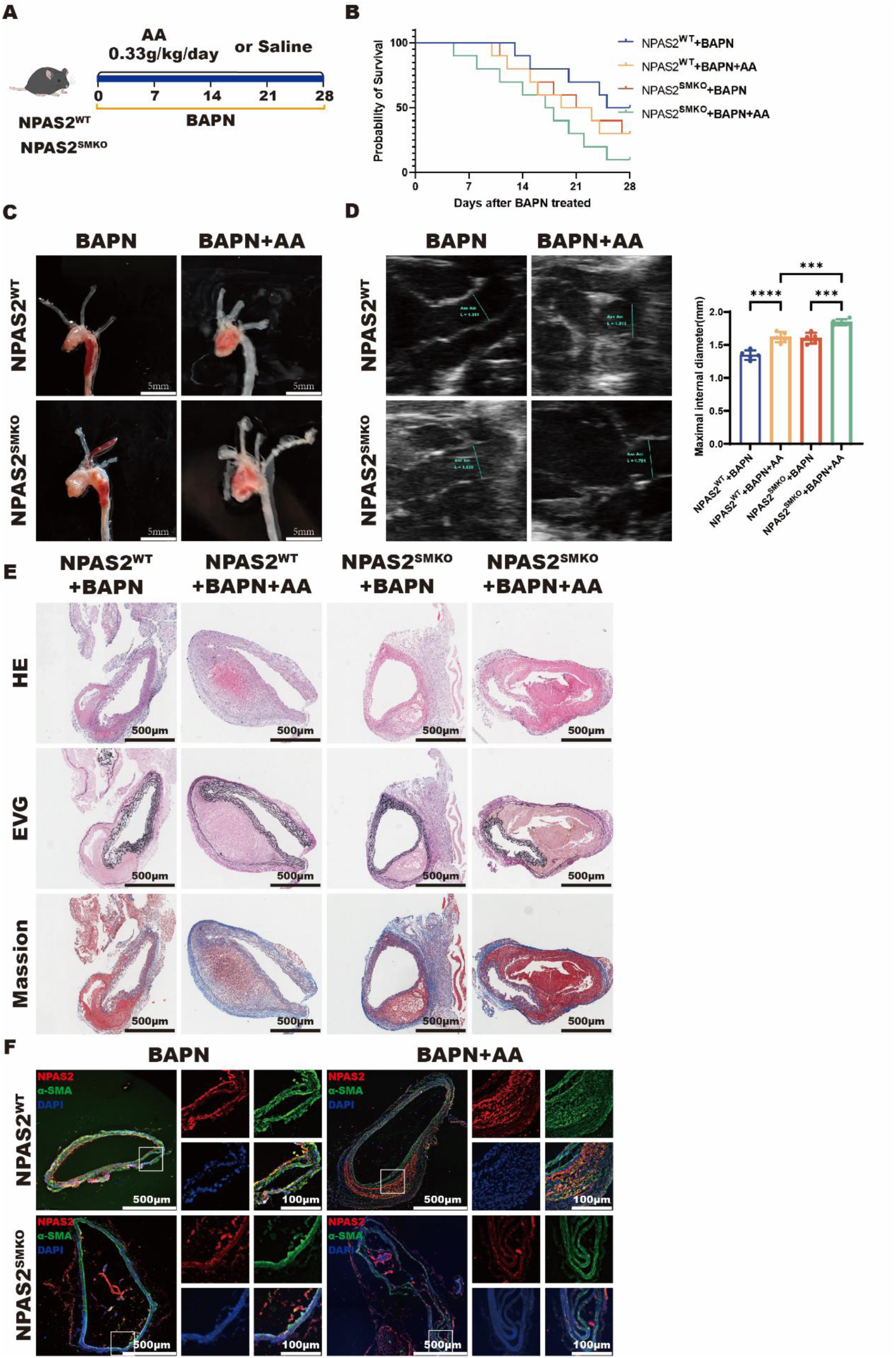
AA aggravates BAPN-induced ATAA in NPAS2^SMKO^ mice. (A)Following a standard diet and BAPN administration, NPAS2^WT^ mice and NPAS2^SMKO^ mice (male, 3 weeks old) were treated with saline or AA for 28 days. n = 10. (B) Survival probability of BAPN-administered NPAS2^WT^ mice and NPAS2^SMKO^ mice treated with saline or AA at 7, 14, 21 and 28 days. n = 10. (C) Representative morphology of ascending thoracic aortas in BAPN-administered NPAS2^WT^ mice and NPAS2^SMKO^ mice treated with saline or AA. Scale bars: 5 mm. n = 10. (D) Representative ultrasound images and inner diameter quantification of ascending thoracic aortas in BAPN-administered NPAS2^WT^ mice and NPAS2^SMKO^ mice treated with saline or AA. n = 8-10. (E) Representative H&E, EVG and Masson staining of ascending thoracic aortas in BAPN-administered NPAS2^WT^ mice and NPAS2^SMKO^ mice treated with saline or AA. Scale bars: 500 μm. n = 6. (F) Representative immunofluorescence images of NPAS2 (red), α-SMA (green), and DAPI (blue) in BAPN-administered NPAS2^WT^ mice and NPAS2^SMKO^ mice treated with saline or AA. Scale bars: 500 μm. n = 6. ***p< 0.001, ****p< 0.0001. B, Kaplan-Meier; D, one-way ANOVA.

### LPCAT3 regulate NPAS2 depletion-induced lipid metabolic disorders

To unravel the potential mechanism underlying NPAS2-regulated lipid signaling networks in phenotypic switching, we performed transcriptome RNA sequencing analysis of HASMCs with and without NPAS2 knockdown to identify the candidate genes, including DPEP1, ACSF2, ACAA1, HSD17B8, LPCAT3, PTGS2, PLCXD2, DHRS9 (Figure 7A). qRT-PCR analysis confirmed LPCAT3 as the most significantly altered gene (Figure 7B). Western blot analysis showed significantly increased LPCAT3 protein levels in NPAS2-knockdown HASMCs (Figure 7C) and in ATAA tissues (Figure 7D). To further substantiate the mechanistic hypothesis, we analyzed lipid profile alterations of HASMCs with or without LPCAT3 overexpression. After transfection with oe-LPCAT3, HASMCs accumulated higher levels of neutral lipids, as demonstrated by BODIPY^493/503^ staining (Figure 7E). As expected, LC-MS/MS analysis of oe-Control and oe-LPCAT3 HASMCs indicated significantly increases in the total lipids (Figure 7F). Further analysis elucidated statistically significant lipid content in oe-Control and oe-LPCAT3 HASMCs (Figure 7G), leading to the identification of eight NPAS2-related lipids (PE, Cer, MG, LPC, FA, DG, PC and PI) (Figure 7H). To assess the impact of LPCAT3 on lipid metabolism, we evaluated fatty acid saturation profiles in oe-LPCAT3 HASMCs by LC-MS/MS. LPCAT3 overexpression increased PUFA content while reducing SFA and MUFA content in PC species. The fatty acid saturation of other phospholipid classes was also significantly modified (Figure 7I). We subsequently investigated LPCAT3-mediated induction of VSMC phenotypic switching. LPCAT3 overexpression elevated OPN and KLF4 protein levels, downregulated α-SMA and SM22-α protein levels (Figure 8A), decreased α-SMA expression (Figure 8B), increased MMP-2 and MMP-9 activities (Figure 8C), and reduced HASMC contractility (Figure 8D).Conversely, LPCAT3 knockout decreased OPN and KLF4 protein levels, enhanced α-SMA and SM22-α protein levels (Figure 8E), augmented α-SMA expression (Figure 8F), diminished MMP-2 and MMP-9 activities (Figure 8G), and improved HASMC contractility (Figure 8H). Moreover, si-LPCAT3 partially reversed si-NPAS2-induced phenotypic switching, confirmed by downregulated synthetic protein levels and upregulated contractile protein levels (Figure 8I), suppressed MMP-2 and MMP-9 activities (Figure 8J), and restored HASMC contractility (Figure 8K). Consequently, NPAS2-mediated transcriptional repression of LPCAT3 inhibits PC-PUFA_2S_-induced VSMC phenotypic switching.

**Figure 7.**
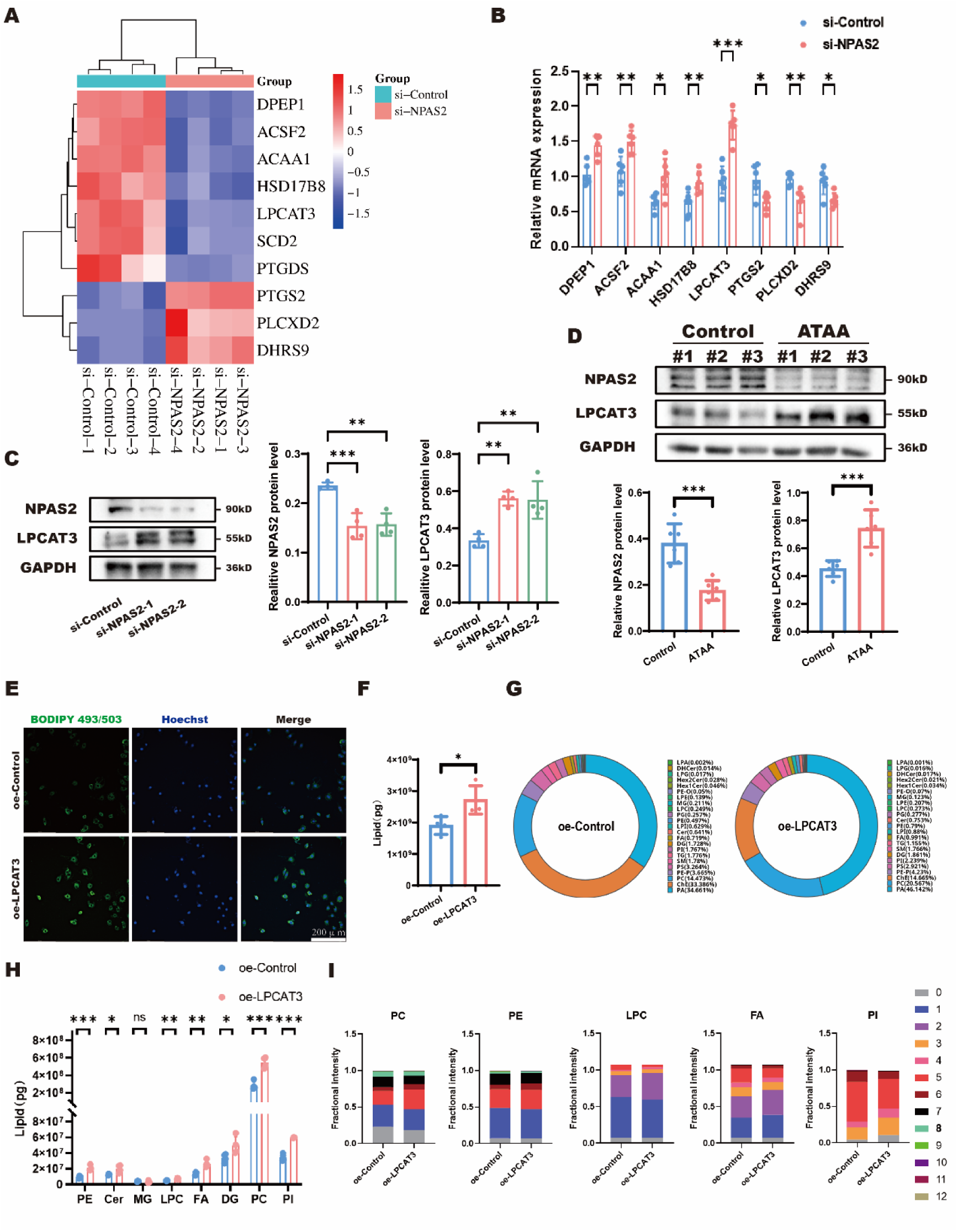
NPAS2 inhibits LPCAT3 expression to regulate lipid metabolism in HASMCs. (A) Transcriptome RNA sequencing analysis of lipid metabolism-related genes in HASMCs transfected with si-Control or si-NPAS2. n = 4. (B) mRNA expression levels of lipid metabolism-related genes (DPEP1, ACSF2, ACAA1, HSD17B8, LPCAT3, PTGS2, PLCXD2, DHRS9) in si-Control and si-NPAS2 HASMCs. n = 6. (C) Representative immunoblots and quantification of NPAS2 and LPCAT3 in si-Control and si-NPAS2 HASMCs. n = 4. (D) Representative immunoblots and quantification of NPAS2 and LPCAT3 in control and ATAA tissues. n = 6. (E) Representative immunofluorescence images of lipid droplets in HASMCs transfected with empty plasmid (oe-Control) or LPCAT3 over-expression plasmid (oe-LPCAT3). Scale bars: 200 μm. n = 6. (F) Lipidomic liquid chromatography–tandem mass spectrometry (LC-MS/MS) analysis of total lipid content in oe-Control and oe-LPCAT3 HASMCs. n = 4. (G) LC-MS/MS analysis of statistically significant lipid species in oe-Control and oe-LPCAT3 HASMCs. n = 4. (H) LC-MS/MS analysis of eight NPAS2-related lipids (PE, Cer, MG, LPC, FA, DG, PC and PI) in oe-Control and oe-LPCAT3 HASMCs. n = 4. (I) Stacked bar charts showing fractional intensity of lipids within lipid classes in oe-Control and oe-LPCAT3 HASMCs. n = 4. *p < 0.05, **p< 0.01, ***p< 0.001. B, F, H and I, unpaired two-tailed t-test; C and D, one-way ANOVA.

**Figure 8.**
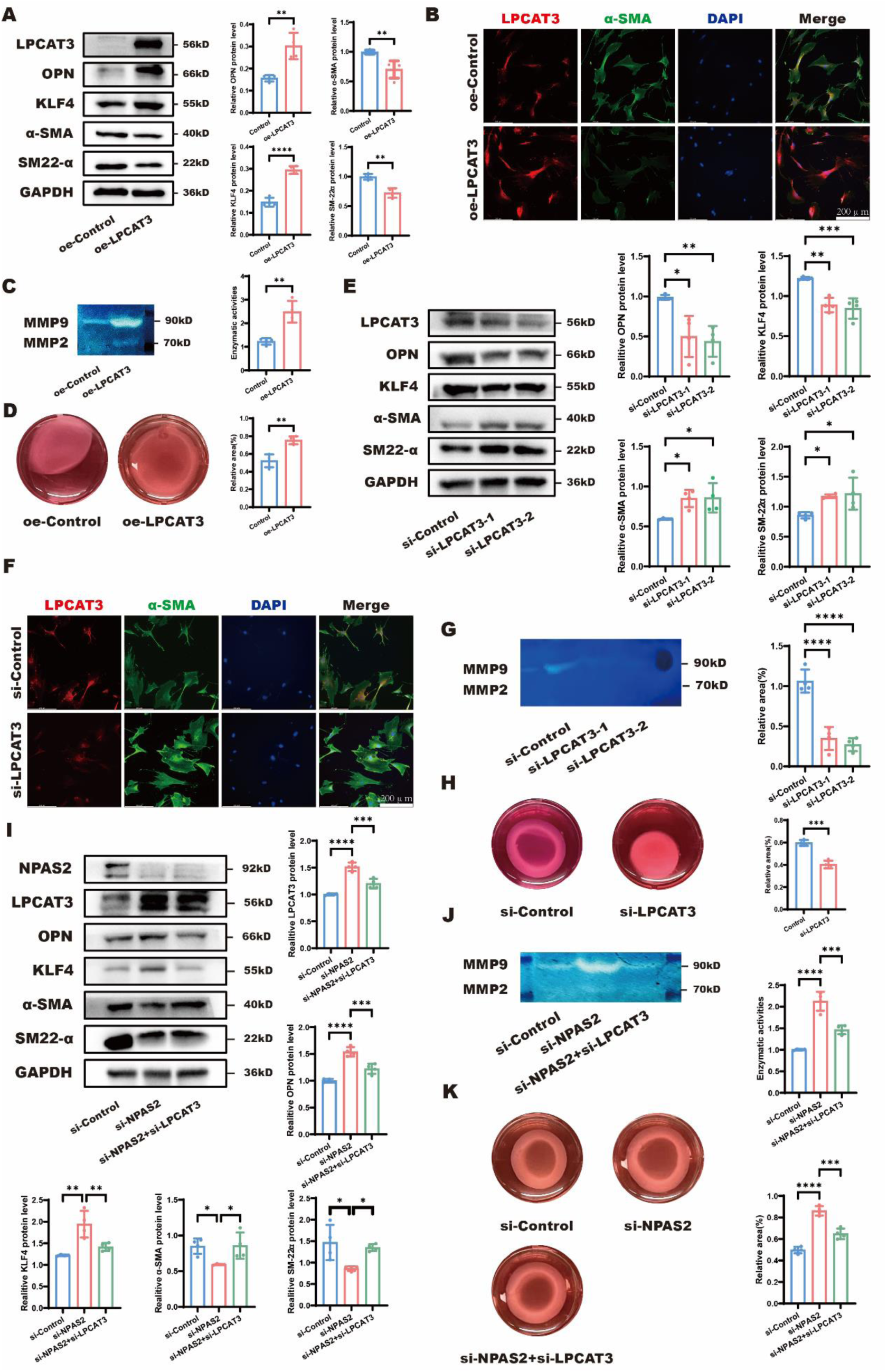
LPCAT3 reverses NPAS2-mediated inhibition of phenotypic switching in HASMCs. (A) Representative immunoblots and quantification of LPCAT3, OPN, KLF4, α-SMA, and SM22-α in HASMCs transfected with empty plasmid (oe-Control) or LPCAT3 over-expression plasmid (oe-LPCAT3). n = 4. (B) Representative immunofluorescence images of LPCAT3 (red), α-SMA (green) and DAPI (blue) in oe-Control and oe-LPCAT3 HASMCs. Scale bars: 200 μm. n = 6. (C) Representative gelatin zymography images and quantification of MMP-2 and MMP-9 enzyme activities in oe-Control and oe-LPCAT3 HASMCs. n = 5. (D) Representative contraction images and quantification of Collagen I in oe-Control and oe-LPCAT3 HASMCs. Scale bars: 5 mm. n = 5. (E) Representative immunoblots and quantification of LPCAT3, OPN, KLF4, α-SMA, and SM22-α in HASMCs transfected with control siRNA (si-Control) or siRNA for LPCAT3 (si-LPCAT3). n = 4. (F) Representative immunofluorescence images of LPCAT3 (red), α-SMA (green) and DAPI (blue) in si-Control and si-LPCAT3 HASMCs. Scale bars: 200 μm. n = 6. (G) Representative gelatin zymography images and quantification of MMP-2 and MMP-9 enzyme activities in si-Control and si-LPCAT3 HASMCs. n = 5. (H) Representative contraction images and quantification of Collagen I in si-Control and si-LPCAT3 HASMCs. Scale bars: 5 mm. n = 5. (I) Representative immunoblots and quantification of NPAS2, LPCAT3, OPN, KLF4, α-SMA, and SM22-α in HASMCs transfected with control siRNA (si-Control), siRNA for NPAS2 (si-NPAS2), siRNA for NPAS2 and LPCAT3 (si-NPAS2+si-LPCAT3). n = 4. (J) Representative gelatin zymography images and quantification of MMP-2 and MMP-9 enzyme activities in HASMCs transfected with si-Control, si-NPAS2 or si-NPAS2+si-LPCAT3. n = 5. (K) Representative contraction images and quantification of Collagen I in HASMCs transfected with si-Control, si-NPAS2 or si-NPAS2+si-LPCAT3. Scale bars: 5 mm. n = 5. *p < 0.05, **p< 0.01, ***p< 0.001, ****p< 0.0001. A, C, D and H, unpaired two-tailed t-test; E, G, I, J and K, one-way ANOVA.

### NPAS2 mediates transcriptional regulation of LPCAT3 by binding to its promoter region

Previous studies have reported that NPAS2 binds to the CYP1A2 promoter in the liver of Npas2^⁻/⁻^ mice⁹. In our study, we observed upregulated LPCAT3 expression in PDGF-BB-treated HASMCs (Figure 9A). Knockdown of NPAS2 further enhanced LPCAT3 transcription both under basal conditions and upon PDGF-BB stimulation (Figure 9B). To explore the underlying mechanism of transcriptional repression, we investigated whether NPAS2 binds to the LPCAT3 promoter using 10 pairs of primers covering various regions (Figure 9C). ChIP assays demonstrated that NPAS2 occupies a region between -550 bp and -250 bp upstream of the LPCAT3 transcription start site (Figure 9C). Both PDGF-BB treatment and NPAS2 knockdown reduced the enrichment of the LPCAT3 promoter fragments immunoprecipitated by an NPAS2 antibody (Figure 9D). Luciferase reporter assays further confirmed that NPAS2 suppresses the transcriptional activity of the LPCAT3 promoter, using a construct containing the -550 bp to -250 bp region (Figure 9E). Importantly, mutation of the predicted binding sites (located between -500 bp and -250 bp) abolished the repressive effect of NPAS2 on the LPCAT3 promoter, highlighting the functional importance of this region in NPAS2-mediated transcriptional regulation (Figure 9E). Together, these findings indicate that NPAS2 inhibits LPCAT3 transcription via binding to LPCAT3’S promoter in HASMCs.

**Figure 9.**
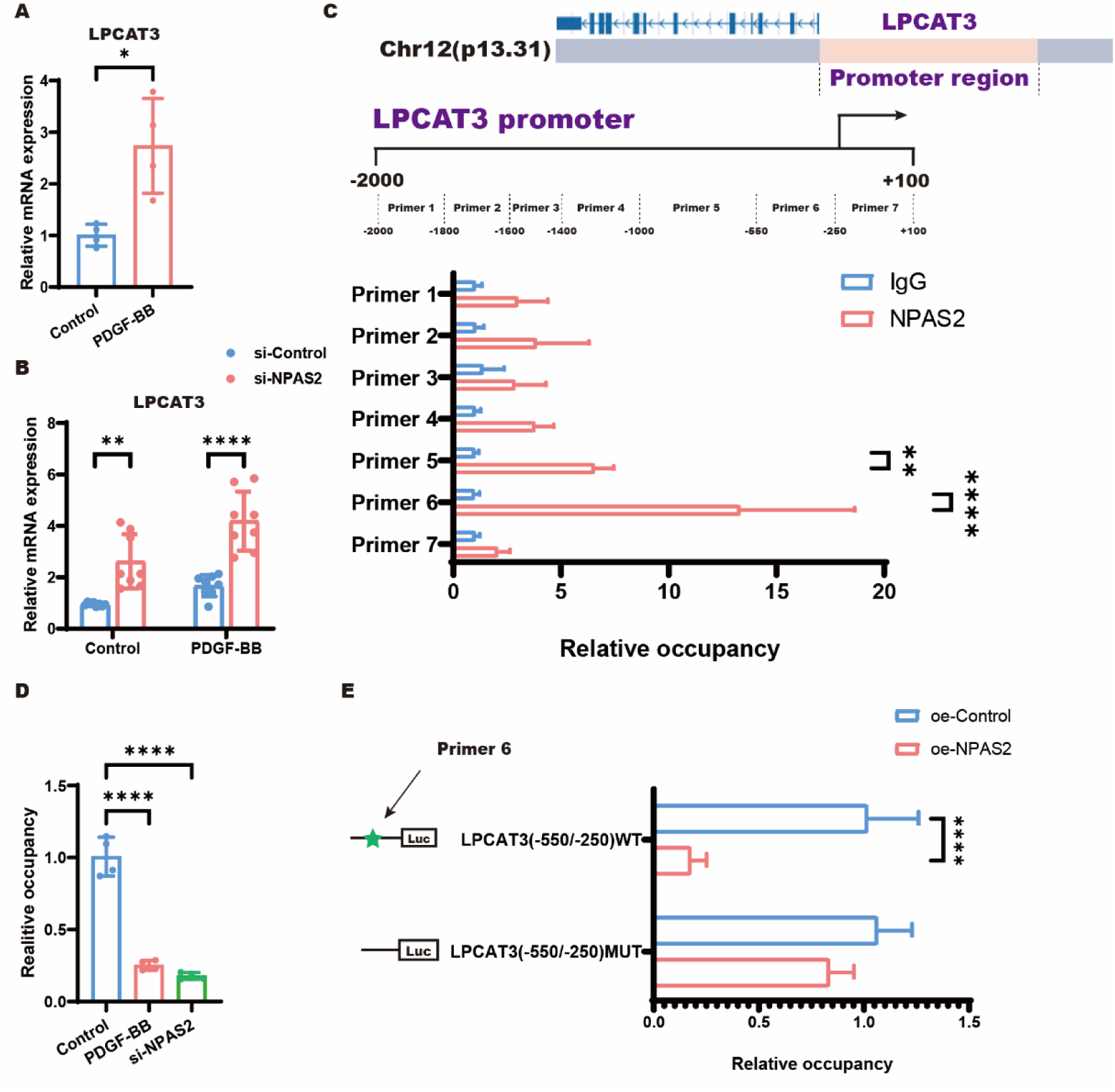
The transcription of LPCAT3 is regulated by NPAS2. (A) Quantification of LPCAT3 mRNA levels in control and PDGF-BB-treated HASMCs (n = 4). (B) LPCAT3 transcriptional levels in HASMCs transfected with si-Control or si-NPAS2, followed by treatment with DMSO or PDGF-BB, as measured by PCR (n = 8). (C) Chromatin immunoprecipitation (ChIP) analysis of NPAS2 binding to the LPCAT3 promoter in HASMCs using an NPAS2-specific antibody or IgG control. NPAS2 occupancy is shown relative to the IgG background signal (n = 4). (D) ChIP analysis of NPAS2 binding to the LPCAT3 promoter in HASMCs treated with Control, PDGF-BB, or si-NPAS2 for 72 hours (n = 4). (E) Relative luciferase activity in HEK293 cells of luciferase reporter constructs containing LPCAT3 promoter truncations or its mutants transfected along with pRL-TK (internal control plasmid) followed by transfection with NPAS2 encoding plasmid. *P < 0.05; **P < 0.01; ****P < 0.0001. A, B, C, E, unpaired two-tailed t-test; D, one-way ANOVA.

### NPAS2-LPCAT3 lipid metabolic axis inhibits ferroptosis to prevent VSMC phenotypic switching

The aforementioned transcriptome RNA sequencing additionally demonstrated statistically significant ferroptosis pathway alterations (Figure 4A). Ferroptosis critically involves biological processes, such as polyunsaturated fatty acid metabolism, iron metabolism, and phospholipids biosynthesis.^9, 10^ We propose ferroptosis represents a nexus integrating NPAS2, lipid metabolism, and phenotypic switching. To confirm NPAS2 involvement in ferroptosis and explore its role in other cell death modalities, we co-treated HASMCs with si-NPAS2 and different cell death inhibitors, including apoptosis inhibitor (Z-VAD-FMK), autophagy inhibitor (bafilomycin A1, Baf-A1), necroptosis inhibitor (necrostatin-1s, Nec-1s), and ferroptosis inhibitor (Ferrostatin-1, Fer-1). Only ferroptosis inhibitors attenuated si-NPAS2-induced cell death (Figure 10A).

**Figure 10.**
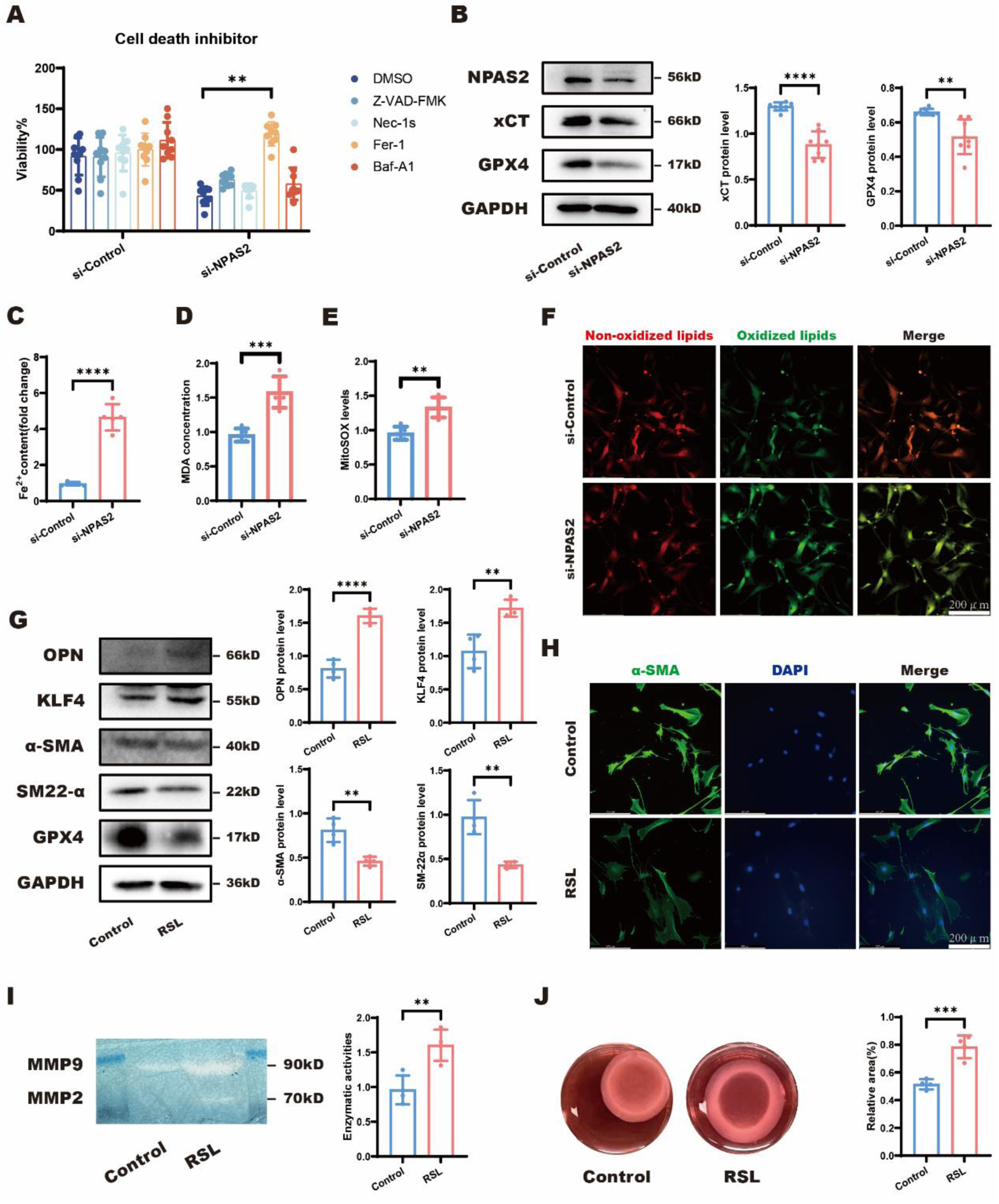
NPAS2 inhibits ferroptosis in HASMCs. (A) Viability of si-Control or si-NPAS2 HASMCs treated with 20 μM Z-VAD-FMK, 20 μM necrostatin-1s (Nec-1s), 10 μM ferrostatin-1 (Fer-1), 100 nM bafilomycin A1 (Baf-A1) for 72 h. n = 6. (B) Representative immunoblots and quantification of NPAS2, xCT and GPX4 in HASMCs transfected with si-Control or si-NPAS2. n = 4. (C) Quantification of Fe^2+^ content in si-Control and si-NPAS2 HASMCs. n = 6. (D) Quantification of MDA concentration in si-Control and si-NPAS2 HASMCs. n = 6. (E) Quantification of MitoSOX levels in si-Control and si-NPAS2 HASMCs. n = 6. (F) Representative immunofluorescence images of non-oxidized lipids (red) and oxidized lipids (green) in si-Control and si-NPAS2 HASMCs. Scale bars: 200 μm. n = 6. (G) Representative immunoblots and quantification of OPN, KLF4, α-SMA, SM22-α, and GPX4 in control and RSL3-treated HASMCs. n = 4. (H) Representative immunofluorescence images of α-SMA (green) and DAPI (blue) in control and RSL3-treated HASMCs. Scale bars: 200 μm. n = 6. (I) Representative gelatin zymography images and quantification of MMP-2 and MMP-9 enzyme activities in control and RSL3-treated HASMCs. n = 5. (J)Representative contraction images and quantification of Collagen I in control and RSL3-treated HASMCs. Scale bars: 5 mm. n = 5. **p< 0.01, ***p< 0.001, ****p< 0.0001. B, C, D, E, G, I and J, unpaired two-tailed t-test; A, one-way ANOVA.

After HASMCs were transfected with si-NPAS2, ferroptosis-related protein levels decreased (xCT and GPX4) (Figure 10B). si-NPAS2-transfected HASMCs simultaneously exhibited increased intracellular Fe^2+^ (Figure 10C), malondialdehyde (MDA; PUFA peroxidation end-product) (Figure 10D), and mitochondrial ROS (mitoSOX) (Figure 10E). BODIPY-C11^581/591^ immunofluorescence detected elevated cellular lipid peroxidation (LPO) in NPAS2-knockdown HASMCs (Figure 10F). To confirm that ferroptosis induces VSMC phenotypic switching, we treated HASMCs with the ferroptosis inducer RSL3.RSL3 triggered phenotypic switching in HASMCs, charactered by upregulated OPN and KLF4 levels alongside downregulated α-SMA, SM22-α and GPX4 levels (Figure 10G), decreased α-SMA expression (Figure 10H), enhanced MMP-2 and MMP-9 activities (Figure 10I), and diminished contractile function (Figure 10J).

We subsequently treated HASMCs with Fer-1 in an attempt to rescue si-NPAS2-induced ferroptosis and attenuate VSMC phenotypic switching.Fer-1 partially rescued both processes in NPAS2-knockdown HASMCs, as demonstrated by downregulated OPN and KLF4 with upregulated α-SMA, SM22-α and XCT (Figure 11A), increased α-SMA expression (Figure 11B), suppressed MMP-2 and MMP-9 activities (Figure 11C), and rescued HASMC contractility (Figure 11D).Thus, ferroptosis facilitates phenotypic switching in si-NPAS2-transfected HASMCs.

**Figure 11.**
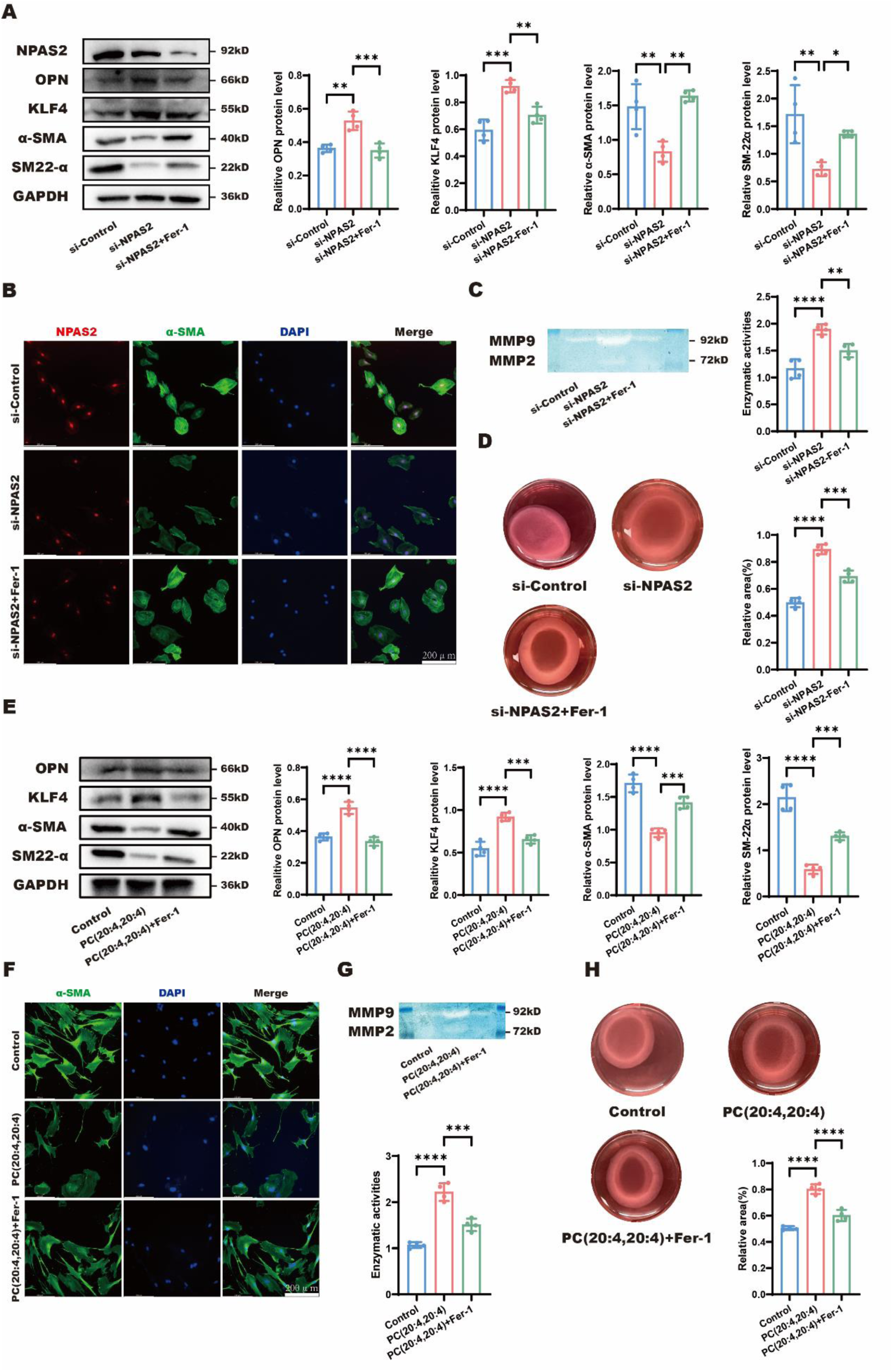
Fer-1 rescues NPAS2-LPCAT3 lipid metabolic axis induced VSMC phenotype switching. (A) Representative immunoblots and quantification of NPAS2, OPN, KLF4, α-SMA and SM22-α in HASMCs treated with control siRNA (si-Control), siRNA for NPAS2 (si-NPAS2), siRNA for NPAS2 and Fer-1 (si-NPAS2 + Fer-1). n = 4. (B) Representative immunofluorescence images of NPAS2 (red), α-SMA (green), and DAPI (blue) in HASMCs treated with si-Control, si-NPAS2, si-NPAS2 + Fer-1. Scale bars: 200 μm. n = 6. (C) Representative gelatin zymography images and quantification of MMP-2 and MMP-9 enzyme activities in HASMCs treated with si-Control, si-NPAS2, si-NPAS2 + Fer-1. n = 5. (D) Representative contraction images and quantification of Collagen I in HASMCs treated with si-Control, si-NPAS2, si-NPAS2 + Fer-1. Scale bars: 5 mm. n = 5. (E) Representative immunoblots and quantification of OPN, KLF4, α-SMA, and SM22-α in HASMCs treated with DMSO, PC (20:4, 20:4), PC (20:4, 20:4) + Fer-1. n = 4. (F) Representative immunofluorescence images of α-SMA (green) and DAPI (blue) in HASMCs treated with DMSO, PC (20:4, 20:4), PC (20:4, 20:4) + Fer-1. Scale bars: 200 μm. n = 6. (G) Representative gelatin zymography images and quantification of MMP-2 and MMP-9 enzyme activities in HASMCs treated with DMSO, PC (20:4, 20:4), PC (20:4, 20:4) + Fer-1. n =5. (H) Representative contraction images and quantification of Collagen I in HASMCs treated with DMSO, PC (20:4, 20:4), PC (20:4, 20:4) + Fer-1. Scale bars: 5 mm. n = 5. * p < 0.05, **p< 0.01, ***p< 0.001, ****p< 0.0001. A, C, D, E, G and H, one-way ANOVA.

To verify the mechanistic hypothesis that PC-PUFA_2S_ contributes to ferroptosis, we treated HASMCs with PC (20:4, 20:4) in the presence or absence of Fer-1 to assess ferroptosis and phenotypic status. As predicted, Fer-1 substantially attenuated PC (20:4, 20:4)-induced ferroptosis to alleviate phenotypic switching, charactered by reduced OPN and KLF4, elevated α-SMA and SM22-α (Figure 11E), augmented α-SMA expression (Figure 11F), inhibited MMP-2 and MMP-9 activities (Figure 11G), and restored HASMC contractility (Figure 11H).

To determine whether NPAS2 deficiency promotes VSMC phenotypic switching through ferroptosis in vivo, we administered Lip-1 to NPAS2 ^WT^ and NPAS2 ^SMKO^ mice with BAPN treatment (Figure 12A). After Lip-1 administration, NPAS2 ^SMKO^ mice with BAPN treatment were less likely to develop ATAA with less ferroptosis degree, evidenced by improved survival probability (Figure 12B), attenuated thoracic aortic dilations/aneurysms (Figure 12C), smaller maximal internal aortic diameters (Figure 12D), slighter disruption of vascular architecture, lower degradation of muscle fibers, less hyperplasia of collagen fibers, reduced fragmentation and degradation of elastic fibers (Figure 12E), and elevated α-SMA expression (Figure 12F).

**Figure 12.**
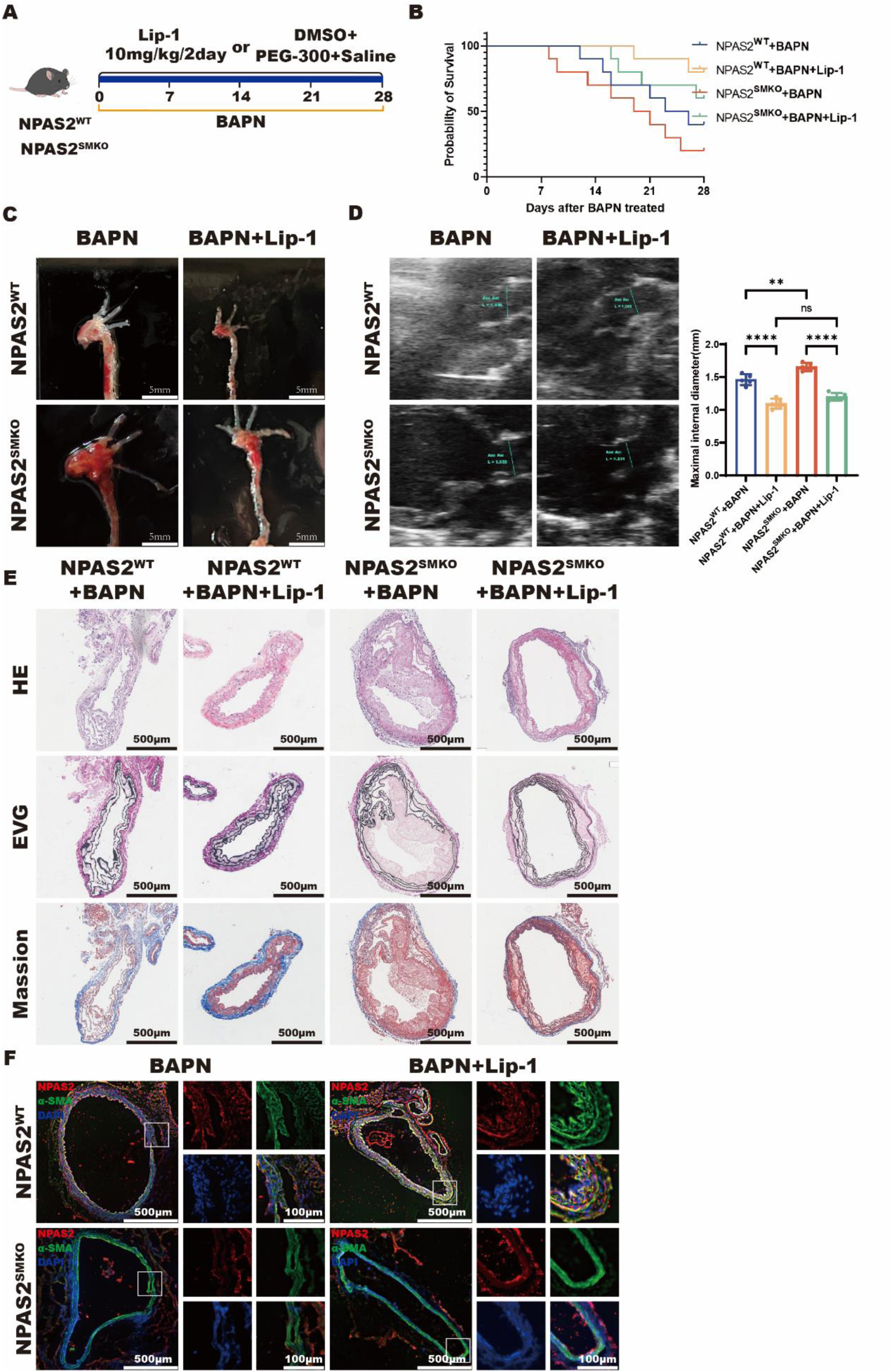
Lip-1 alleviates BAPN-induced ATAA in NPAS2^SMKO^ mice. (A) Following a standard diet and BAPN administration, NPAS2^WT^ mice and NPAS2^SMKO^ mice (male, 3 weeks old) were treated with saline+ DMSO+ PEG-300 or Lip-1 for 28 days. n = 10. (B) Survival probability of BAPN-administered NPAS2^WT^ mice and NPAS2^SMKO^ mice treated with saline+ DMSO+ PEG-300 or Lip-1 at 7, 14, 21 and 28 days. n = 10. (C) Representative morphology of ascending thoracic aortas in BAPN-administered NPAS2^WT^ mice and NPAS2^SMKO^ mice treated with saline+ DMSO+ PEG-300 or Lip-1. Scale bars: 5 mm. n = 10. (D) Representative ultrasound images and inner diameter quantification of ascending thoracic aortas in BAPN-administered NPAS2^WT^ mice and NPAS2^SMKO^ mice treated with saline+ DMSO+ PEG-300 or Lip-1. n = 10. (E) Representative H&E, EVG and Masson staining of ascending thoracic aortas in BAPN-administered NPAS2^WT^ mice and NPAS2^SMKO^ mice treated with saline+ DMSO+ PEG-300 or Lip-1. Scale bars: 500 μm. n = 6. (F) Immunofluorescence of NPAS2 (red), α-SMA (green), and DAPI (blue) in BAPN-administered NPAS2^WT^ mice and NPAS2^SMKO^ mice treated with saline+ DMSO+ PEG-300 or Lip-1. Scale bars: 500 μm. n = 6. **p< 0.01, ****p< 0.0001. B, Kaplan-Meier; D, one-way ANOVA.

NPAS2 deficiency facilitates PC-PUFA_2S_-induced ferroptosis, ultimately exacerbating the VSMC phenotypic switching in ATAA.

## Discussion

NPAS2, a core circadian transcription factor, has been extensively studied in forebrain and suprachiasmatic nucleus, while its peripheral functions remain largely uncharacterized. Previous studies demonstrated that NPAS2 critically regulated hepatic lipid metabolism, with Npas2^-/-^ mice exhibiting significant alterations in fatty acid metabolism pathways^7, 11^ Mechanistically, NPAS2 interacted with the hepatic small heterodimer partner (SHP) to establish a feedback regulatory loop maintaining hepatic lipid homeostasis.^8^ Notably, NPAS2 deficiency triggered profound hepatic steatosis in Shp^−/−^ mice, associated with disrupted triglyceride and lipoprotein metabolism.^12^ NPAS2 activated CYP1A2 transcription through direct promoter binding in the liver of Npas2^−/−^ mice.^13^ CYP1A2, a major form of hepatic cytochrome P450 (CYP) superfamily, mediated the biotransformation of various endogenous compounds including PUFAs and steroid hormones (estradiol, estrone).^14, 15^ Additionally, NPAS2 dysregulation further contributed to alcohol-induced hepatic steatosis.^16^ NPAS2 polymorphisms associated with common metabolic syndrome risk factors. NPAS2 disturbance could lead to hypertension.^17^ The aforementioned studies established NPAS2 as a critical regulator of lipid homeostasis.^12^ Our transcriptomic analysis of human aortic specimens revealed significant NPAS2 alterations in ATAA patients versus controls. In NPAS2-knockdown HASMCs, primary dysregulation mainly focused on fatty acid metabolism (a subset of the lipid metabolism). Thus, NPAS2 is corelated closely with ATAA progression and lipid metabolism. Further investigation is needed to elucidate the detailed mechanism of NPAS2-mediated lipid metabolism in ATAA.

Lipids maintain cellular homeostasis by serving as energetic substrates, membrane components, signaling mediators and protein modifiers.^18, 19^

Lipids are broadly classified into fatty acids, neutral lipids, and phospholipids.^20, 21^ Fatty acids (FAs), core constituents of all lipids, act as biosynthetic precursors for other lipids.^20, 22^ Their intracellular functions dependent not only on the total abundance but also on chain length and structural features (e.g., carbon-carbon double bonds).^23^ Specifically, saturated fatty acids (SFA), mono fatty acids (MUFA), and polyunsaturated fatty acids (PUFA) are in a tightly regulated dynamic balance to maintain intracellular homeostasis and biological functions.^23, 24^ ROS-mediated PUFA peroxidation generates PUFA hydroperoxides (PUFA-OOH), destabilizing membrane architecture, causing oxidative damage to DNA and proteins, and inducing ferroptosis.^25, 26^ Further LC-MS/MS analysis identified statistically significant lipids. Subsequent exogenous lipid treatment identified phosphatidylcholines (PCs) as the most potent phospholipid inducing phenotypic switching in HASMCs. Critically, PCs with two polyunsaturated fatty acyl chains (PC-PUFA_2S_) exhibited markedly stronger induction of VSMC phenotypic switching compared to PCs with one polyunsaturated fatty acyl chain (PC-PUFA_1S_). Decreasing PC-PUFA_2S_ concentrations in humans would represent an important strategy for preventing aortic aneurysm formation.

Acyl-CoA synthetase long-chain family member 4 (ACSL4) and lysophosphatidylcholine acyltransferase 3 (LPCAT3) were characterized as essential enzymes incorporating PUFAs into phospholipids, forming PL-PUFAs and subsequently inducing ferroptosis.^25, 27, 28^ ACSL4 catalyzes the conversion of free PUFAs to PUFA-CoA derivatives (PUFA-CoAs).^25^ LPCAT3 subsequently esterifies PUFA-CoAs into PC-PUFAs, which integrate into plasma membranes and enhance susceptibility to lipid peroxidation.^25, 28^ Peroxidized lipids destabilize the membrane bilayer, making it particularly prone to disruption and ferroptosis.^23, 29^ To unravel the NPAS2-regulated lipid signaling in VSMC phenotypic switching, transcriptome RNA sequencing analysis of NPAS2-knockdown and control HASMCs identified eight significant candidates. qRT-PCR confirmed LPCAT3 as the most profoundly dysregulated gene. Subsequent LC-MS/MS analysis of LPCAT3-overexpressing and control HASMCs indicated eight NPAS2-related lipids. Further, we performed loss-of-function and gain-of-function genetic experiments. Thus, NPAS2 depletion facilitated PC-PUFA_2S_-induced VSMC phenotypic switching through LPCAT3, ultimately exacerbating ATAA pathogenesis. Inhibiting LPCAT3 activity through drug development may provide a novel therapeutic intervention to mitigate disease progression in aortic aneurysm patients.

Regulated cell death (RCD) encompasses spontaneous and programmed cell death in physiological and pathological contexts.^2, 30^ This study focused on well-characterized RCD forms in aortic aneurysm and dissection^2^, including apoptosis, autophagy-dependent cell death, ferroptosis and necroptosis, to determine the specific RCD modality involved in NPAS2-regulated VSMC phenotypic transformation in ATAA. Ferroptosis, a newly identified regulated cell death, is mainly driven by iron-dependent lipid peroxidation.^23, 31^ Ferroptosis has emerged as a significant regulatory mechanism involved in the development of various cardiovascular disorders, such as arrhythmias, myocardial ischemia-reperfusion injury, diabetic cardiomyopathy and atherosclerosis^32^. In this study, transcriptome sequencing further revealed significant ferroptosis pathway dysregulation. We confirmed that NPAS2 specifically modulated ferroptosis, with no significant association with other RCD types. Ferroptosis inhibitor rescued both si-NPAS2-induced and NPAS2^SMKO^-induced ferroptosis, attenuating phenotypic switching in vitro and in vivo. Ferroptosis is closely correlated with VSMC phenotypic switching. Emerging evidence indicates that ferroptosis-associated genes exhibit profound dysregulation in ascending thoracic aortic aneurysm and dissection (ATAAD) tissues compared to normal aortic tissues.^2, 32, 33^ Lipid peroxidation and iron overload, distinguishing characteristics of ferroptosis, contribute to progression of ATAAD. Previous study demonstrated markedly elevated lipid peroxidation levels in the aortic media of ATAAD patients versus normal controls.^2, 34^ In this study, ferroptosis inhibitor considerably rescued PC-PUFA_2S_-induced ferroptosis, thereby alleviating VSMC phenotype switching. Collectively, these findings confirmed the involvement of NPAS2-mediated ferroptosis in ATAA pathogenesis.

Therefore, NPAS2-mediated transcriptional repression of LPCAT3 inhibits PC-PUFA_2S_-induced VSMC phenotypic switching. Clinically, NPAS2 serves as a predictive biomarker for risk stratification and represents a promising therapeutic target.

## Conclusions

NPAS2 can transcriptionally repress LPCAT3 to protect against PC-PUFA_2S_-induced ferroptosis, thereby preventing VSMC phenotypic switching in ATAA. NPAS2 represents a promising therapeutic target for ATAA prevention and treatment.

## Acknowledgements

Not applicable.

## Supplementary data

Supplementary data are available at European Heart Journal online.

## Declarations

### Supplemental Material

supplementary file

### Data availability

Data will be made available on request.

### Ethical Approval

This study involving human participants was approved by the Ethics Committee of the First Affiliated Hospital of Nanjing Medical University (IRB number:2019-SR-067). All procedures complied with the ethical standards stated in the Declaration of Helsinki. Written informed consent was obtained from all participants preoperatively.

This study involving mice was approved by the Nanjing Medical University Animal Care and Use Committee. All animal care and experimental protocols were approved by the Animal Research: Reporting of In Vivo Experiments (ARRIVE) guidelines.

### Pre-registered Clinical Trial Number

Not applicable.

## Author Contributions

Wenfeng Lin: Conceptualization, Methodology, Formal analysis, Investigation. Jiaqi Xiong: Validation, Investigation, Writing - Original Draft. Yuhan Yan: Investigation, Resources, Writing - Review & Editing. Jinhui Bian: Methodology. Zhen Zhang: Resources. Xinyang Xu: Formal analysis. Yifei Diao: Data Curation. Xiaoxia Lan: Software. Yu Zhong: Data Curation. Haibin Ji: Visualization. Liping Xie: Supervision. Yongfeng Shao: Project administration. Buqing Ni: Funding acquisition.

## Supplementary material

### Supplemental Tables

**Supplemental Table S1:**
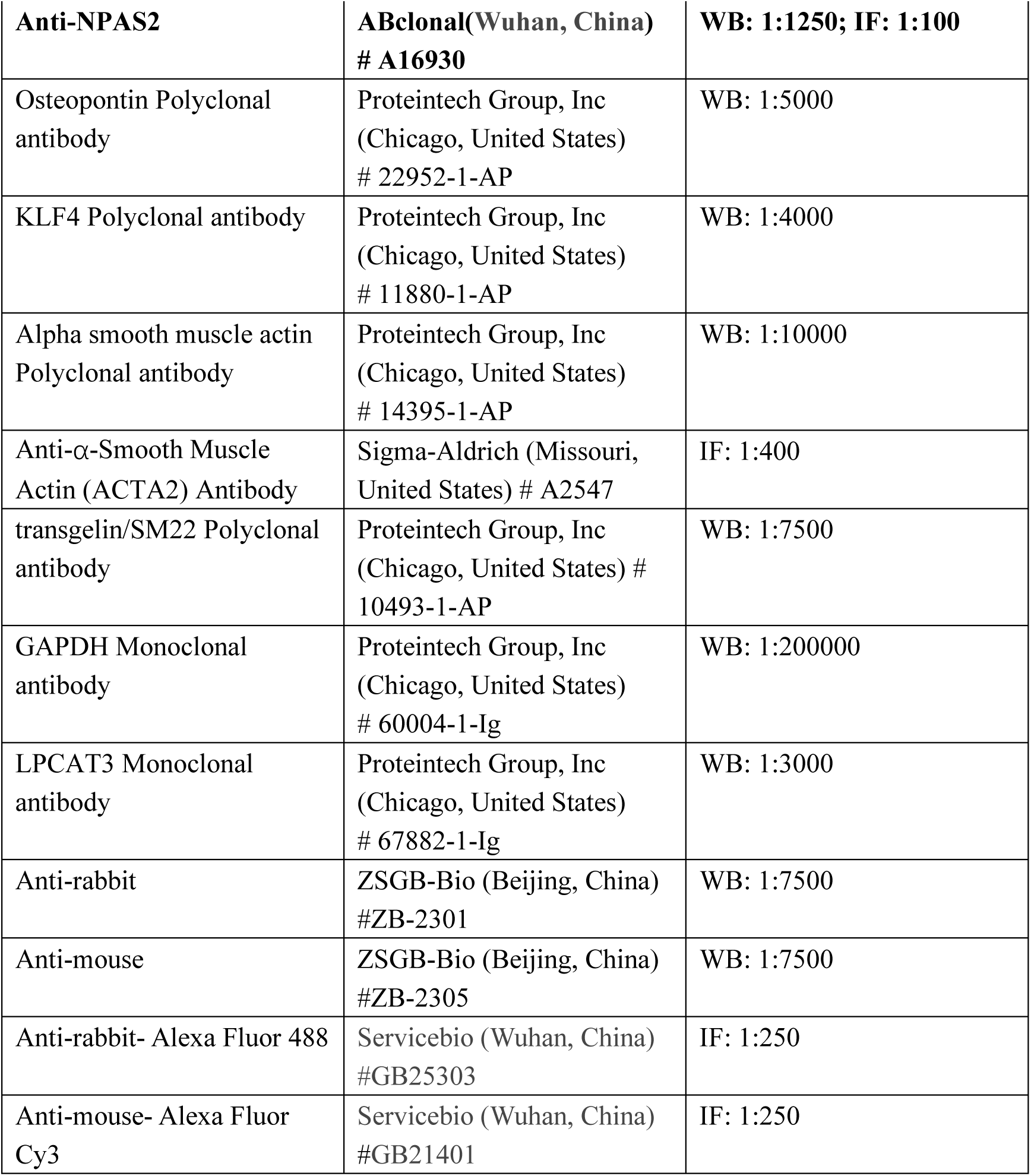
The Antibodies used in this study.

**Supplemental Table S2:**
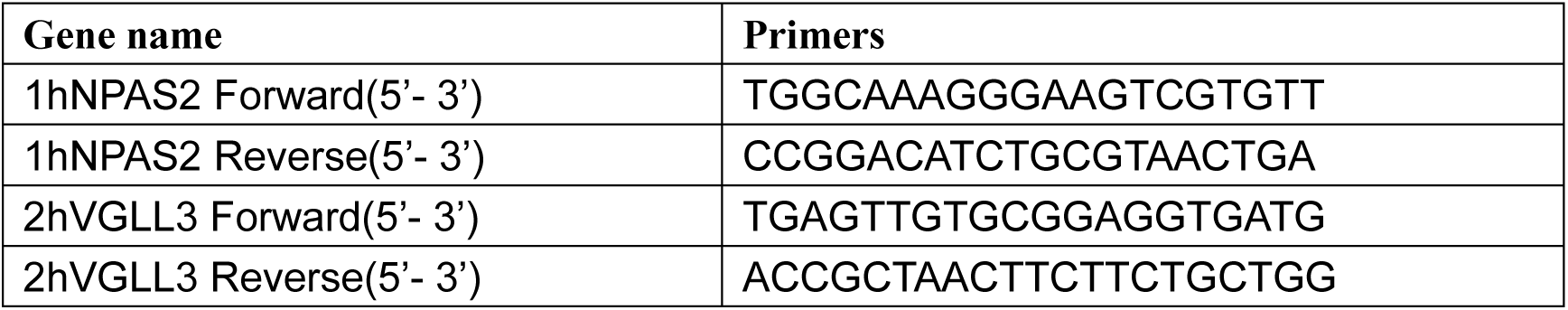
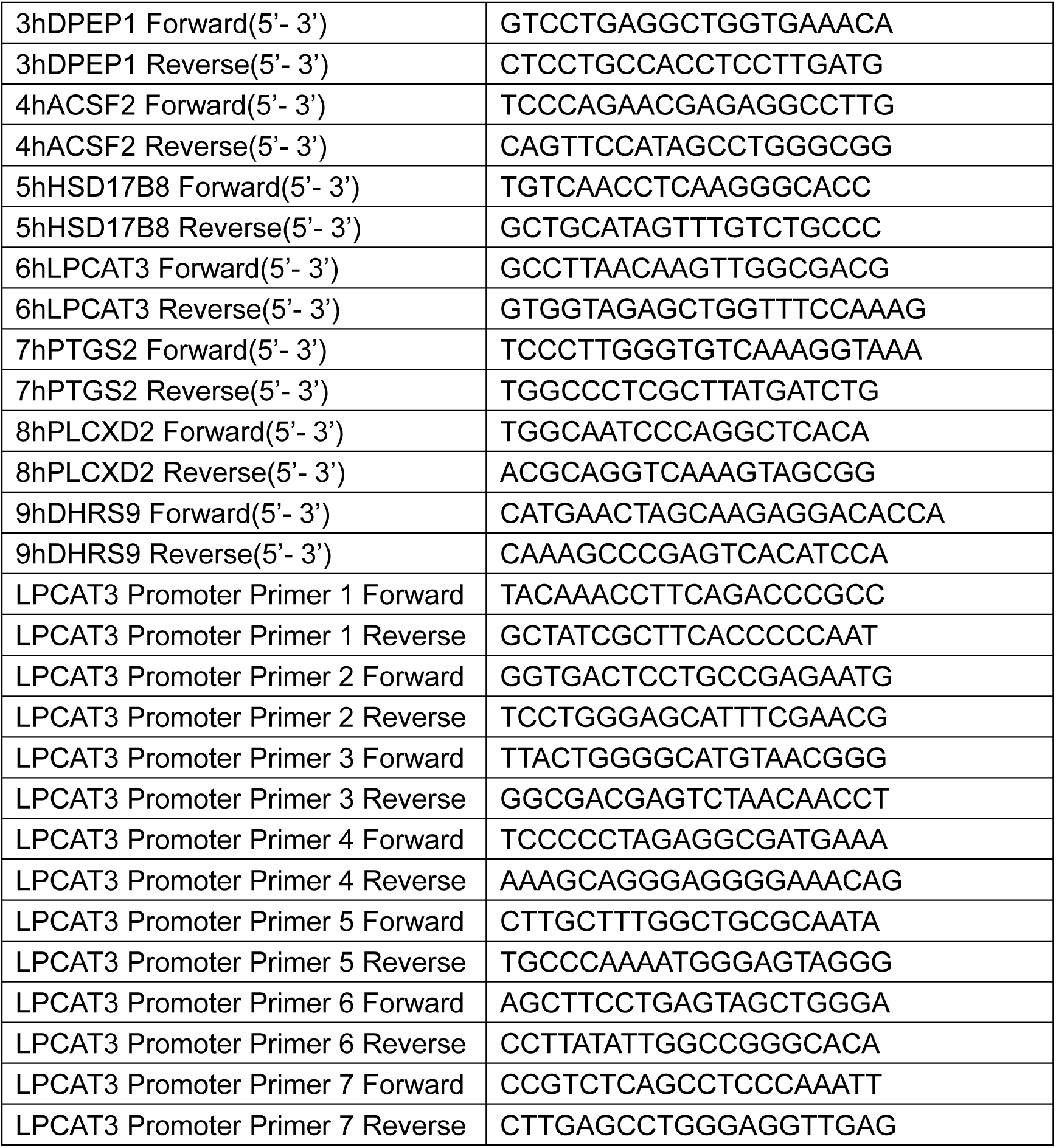
The Primers of Relative Genes in this study.

**Supplemental Table S3:**
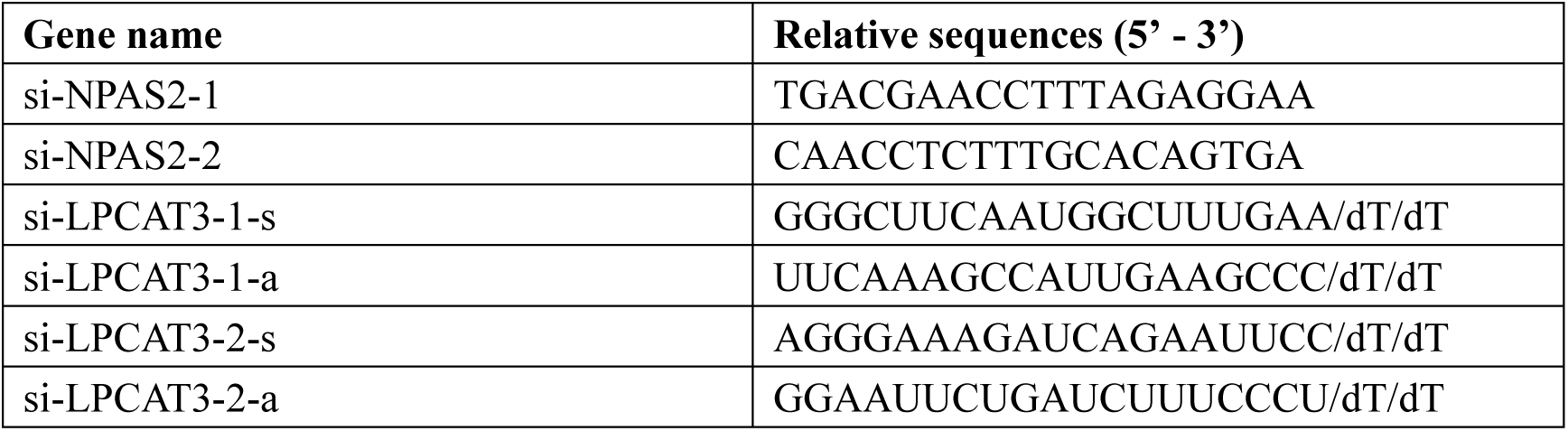
The Relative siRNA in this study.

**Supplemental Figure S1:**
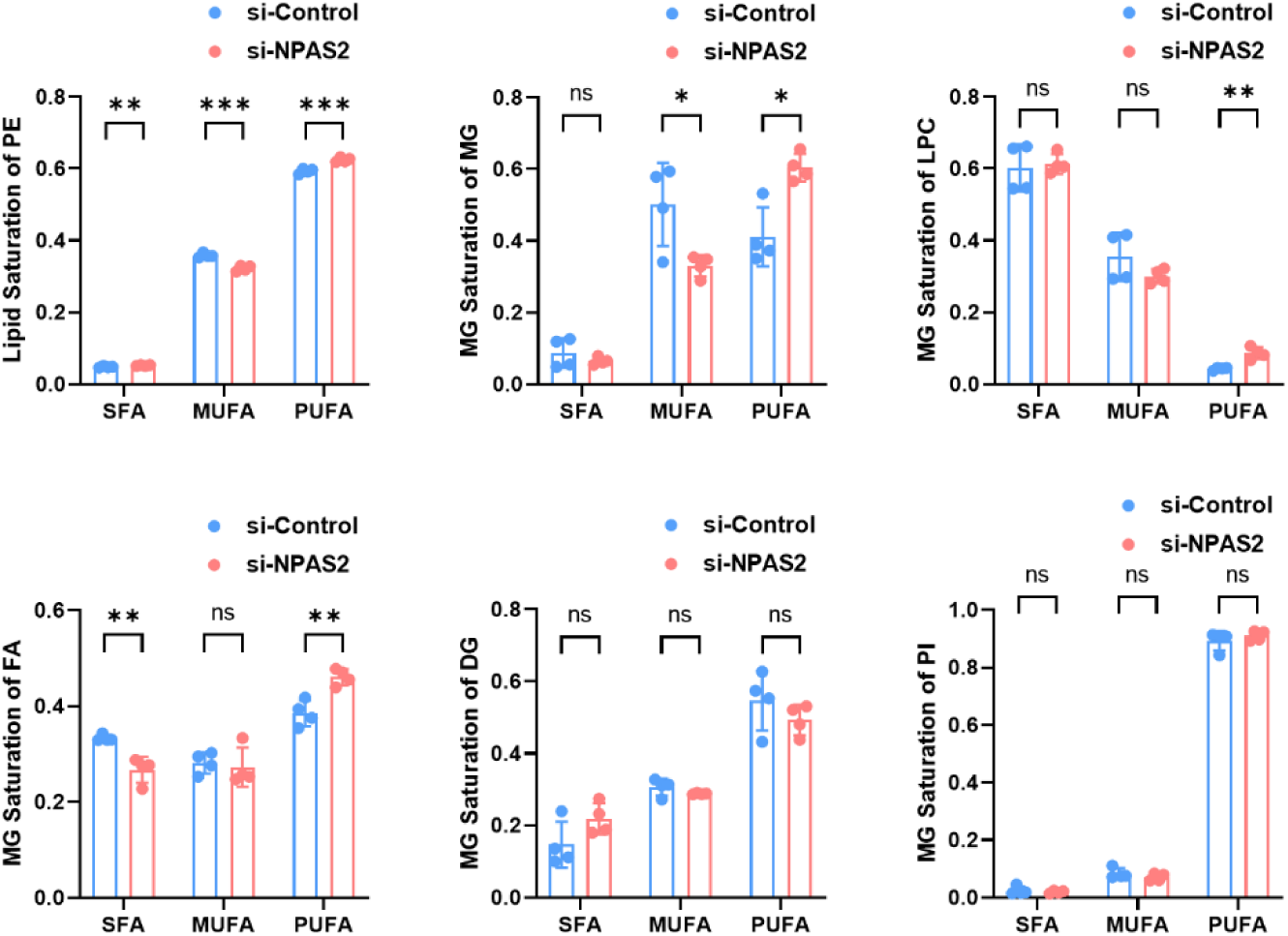
The effect of NPAS2 depletion on fatty acid saturation profiles.

## References

1. Ganizada BH, Veltrop RJA, Akbulut AC, Koenen RR, Accord R, Lorusso R, et al. Unveiling cellular and molecular aspects of ascending thoracic aortic aneurysms and dissections. Basic Res Cardiol. 2024;119(3):371–95.

2. Chen Y, He Y, Wei X, Jiang DS. Targeting regulated cell death in aortic aneurysm and dissection therapy. Pharmacol Res. 2022;176:106048.

3. Cho MJ, Lee MR, Park JG. Aortic aneurysms: current pathogenesis and therapeutic targets. Exp Mol Med. 2023;55(12):2519–30.

4. Song T, Zhao S, Luo S, Chen C, Liu X, Wu X, et al. SLC44A2 regulates vascular smooth muscle cell phenotypic switching and aortic aneurysm. J Clin Invest. 2024;134(16).

5. Qiu Z, Ming H, Lei S, Zhou B, Zhao B, Yu Y, et al. Roles of HDAC3-orchestrated circadian clock gene oscillations in diabetic rats following myocardial ischaemia/reperfusion injury. Cell Death Dis. 2021;12(1):43.

6. Dioum EM, Rutter J, Tuckerman JR, Gonzalez G, Gilles-Gonzalez MA, McKnight SL. NPAS2: a gas-responsive transcription factor. Science. 2002;298(5602):2385–7.

7. Matsumura R, Matsubara C, Node K, Takumi T, Akashi M. Nuclear receptor-mediated cell-autonomous oscillatory expression of the circadian transcription factor, neuronal PAS domain protein 2 (NPAS2). J Biol Chem. 2013;288(51):36548–53.

8. Murgo E, Colangelo T, Bellet MM, Malatesta F, Mazzoccoli G. Role of the Circadian Gas-Responsive Hemeprotein NPAS2 in Physiology and Pathology. Biology (Basel). 2023;12(10).

9. Zhang S, Bei Y, Huang Y, Huang Y, Hou L, Zheng XL, et al. Induction of ferroptosis promotes vascular smooth muscle cell phenotypic switching and aggravates neointimal hyperplasia in mice. Mol Med. 2022;28(1):121.

10. Yang WS, Stockwell BR. Ferroptosis: Death by Lipid Peroxidation. Trends Cell Biol. 2016;26(3):165–76.

11. O’Neil D, Mendez-Figueroa H, Mistretta TA, Su C, Lane RH, Aagaard KM. Dysregulation of Npas2 leads to altered metabolic pathways in a murine knockout model. Mol Genet Metab. 2013;110(3):378–87.

12. Lee SM, Zhang Y, Tsuchiya H, Smalling R, Jetten AM, Wang L. Small heterodimer partner/neuronal PAS domain protein 2 axis regulates the oscillation of liver lipid metabolism. Hepatology. 2015;61(2):497–505.

13. He Y, Cen H, Guo L, Zhang T, Yang Y, Dong D, et al. Circadian oscillator NPAS2 regulates diurnal expression and activity of CYP1A2 in mouse liver. Biochem Pharmacol. 2022;206:115345.

14. Pu X, Gao Y, Li R, Li W, Tian Y, Zhang Z, et al. Biomarker Discovery for Cytochrome P450 1A2 Activity Assessment in Rats, Based on Metabolomics. Metabolites. 2019;9(4).

15. Zhou SF, Liu JP, Chowbay B. Polymorphism of human cytochrome P450 enzymes and its clinical impact. Drug Metab Rev. 2009;41(2):89–295.

16. Zhou P, Ross RA, Pywell CM, Liangpunsakul S, Duffield GE. Disturbances in the murine hepatic circadian clock in alcohol-induced hepatic steatosis. Sci Rep. 2014;4:3725.

17. Englund A, Kovanen L, Saarikoski ST, Haukka J, Reunanen A, Aromaa A, et al. NPAS2 and PER2 are linked to risk factors of the metabolic syndrome. J Circadian Rhythms. 2009;7:5.

18. Fanning S, Haque A, Imberdis T, Baru V, Barrasa MI, Nuber S, et al. Lipidomic Analysis of alpha-Synuclein Neurotoxicity Identifies Stearoyl CoA Desaturase as a Target for Parkinson Treatment. Mol Cell. 2019;73(5):1001–14 e8.

19. Sastry PS. Lipids of nervous tissue: composition and metabolism. Prog Lipid Res. 1985;24(2):69–176.

20. Yoon H, Shaw JL, Haigis MC, Greka A. Lipid metabolism in sickness and in health: Emerging regulators of lipotoxicity. Mol Cell. 2021;81(18):3708–30.

21. Mutlu AS, Duffy J, Wang MC. Lipid metabolism and lipid signals in aging and longevity. Dev Cell. 2021;56(10):1394–407.

22. de Carvalho C, Caramujo MJ. The Various Roles of Fatty Acids. Molecules. 2018;23(10).

23. Lorito N, Subbiani A, Smiriglia A, Bacci M, Bonechi F, Tronci L, et al. FADS1/2 control lipid metabolism and ferroptosis susceptibility in triple-negative breast cancer. EMBO Mol Med. 2024;16(7):1533–59.

24. Dyall SC, Balas L, Bazan NG, Brenna JT, Chiang N, da Costa Souza F, et al. Polyunsaturated fatty acids and fatty acid-derived lipid mediators: Recent advances in the understanding of their biosynthesis, structures, and functions. Prog Lipid Res. 2022;86:101165.

25. Ru Q, Li Y, Chen L, Wu Y, Min J, Wang F. Iron homeostasis and ferroptosis in human diseases: mechanisms and therapeutic prospects. Signal Transduct Target Ther. 2024;9(1):271.

26. Minotti G, Aust SD. The role of iron in oxygen radical mediated lipid peroxidation. Chem Biol Interact. 1989;71(1):1–19.

27. Sun S, Shen J, Jiang J, Wang F, Min J. Targeting ferroptosis opens new avenues for the development of novel therapeutics. Signal Transduct Target Ther. 2023;8(1):372.

28. Doll S, Proneth B, Tyurina YY, Panzilius E, Kobayashi S, Ingold I, et al. ACSL4 dictates ferroptosis sensitivity by shaping cellular lipid composition. Nat Chem Biol. 2017;13(1):91–8.

29. Dixon SJ, Lemberg KM, Lamprecht MR, Skouta R, Zaitsev EM, Gleason CE, et al. Ferroptosis: an iron-dependent form of nonapoptotic cell death. Cell. 2012;149(5):1060–72.

30. Hotchkiss RS, Strasser A, McDunn JE, Swanson PE. Cell death. N Engl J Med. 2009;361(16):1570–83.

31. Stockwell BR, Jiang X, Gu W. Emerging Mechanisms and Disease Relevance of Ferroptosis. Trends Cell Biol. 2020;30(6):478–90.

32. Fang X, Ardehali H, Min J, Wang F. The molecular and metabolic landscape of iron and ferroptosis in cardiovascular disease. Nat Rev Cardiol. 2023;20(1):7–23.

33. Zou HX, Qiu BQ, Lai SQ, Huang H, Zhou XL, Gong CW, et al. Role of ferroptosis-related genes in Stanford type a aortic dissection and identification of key genes: new insights from bioinformatic analysis. Bioengineered. 2021;12(2):9976–90.

34. Liao M, Liu Z, Bao J, Zhao Z, Hu J, Feng X, et al. A proteomic study of the aortic media in human thoracic aortic dissection: implication for oxidative stress. J Thorac Cardiovasc Surg. 2008;136(1):65–72, e1-3.

