## supplementary file for "NPAS2 attenuates VSMC phenotypic switching in ascending thoracic aortic aneurysm via LPCAT3/PC-PUFA2S-mediated ferroptosis"

**Supplementary material**

**Supplemental Tables**

**Supplemental Table S1:** **The Antibodies used in this study**

| **Anti-NPAS2** | **ABclonal(Wuhan, China) # A16930** | **WB: 1:1250; IF: 1:100** |
| --- | --- | --- |
| Osteopontin Polyclonal antibody | Proteintech Group, Inc (Chicago, United States)  # 22952-1-AP | WB: 1:5000 |
| KLF4 Polyclonal antibody | Proteintech Group, Inc (Chicago, United States)  # 11880-1-AP | WB: 1:4000 |
| Alpha smooth muscle actin Polyclonal antibody | Proteintech Group, Inc (Chicago, United States)  # 14395-1-AP | WB: 1:10000 |
| Anti-α-Smooth Muscle Actin (ACTA2) Antibody | Sigma-Aldrich (Missouri, United States) # A2547 | IF: 1:400 |
| transgelin/SM22 Polyclonal antibody | Proteintech Group, Inc (Chicago, United States) # 10493-1-AP | WB: 1:7500 |
| GAPDH Monoclonal antibody | Proteintech Group, Inc (Chicago, United States)  # 60004-1-Ig | WB: 1:200000 |
| LPCAT3 Monoclonal antibody | Proteintech Group, Inc (Chicago, United States)  # 67882-1-Ig | WB: 1:3000 |
| Anti-rabbit | ZSGB-Bio (Beijing, China) #ZB-2301 | WB: 1:7500 |
| Anti-mouse | ZSGB-Bio (Beijing, China) #ZB-2305 | WB: 1:7500 |
| Anti-rabbit- Alexa Fluor 488 | Servicebio (Wuhan, China) #GB25303 | IF: 1:250 |
| Anti-mouse- Alexa Fluor Cy3 | Servicebio (Wuhan, China) #GB21401 | IF: 1:250 |

**Supplemental Table S2:** **The Primers of Relative Genes in this study**

| **Gene name** | **Primers** |
| --- | --- |
| 1hNPAS2 Forward(5’- 3’) | TGGCAAAGGGAAGTCGTGTT |
| 1hNPAS2 Reverse(5’- 3’) | CCGGACATCTGCGTAACTGA |
| 2hVGLL3 Forward(5’- 3’) | TGAGTTGTGCGGAGGTGATG |
| 2hVGLL3 Reverse(5’- 3’) | ACCGCTAACTTCTTCTGCTGG |
| 3hDPEP1 Forward(5’- 3’) | GTCCTGAGGCTGGTGAAACA |
| 3hDPEP1 Reverse(5’- 3’) | CTCCTGCCACCTCCTTGATG |
| 4hACSF2 Forward(5’- 3’) | TCCCAGAACGAGAGGCCTTG |
| 4hACSF2 Reverse(5’- 3’) | CAGTTCCATAGCCTGGGCGG |
| 5hHSD17B8 Forward(5’- 3’) | TGTCAACCTCAAGGGCACC |
| 5hHSD17B8 Reverse(5’- 3’) | GCTGCATAGTTTGTCTGCCC |
| 6hLPCAT3 Forward(5’- 3’) | GCCTTAACAAGTTGGCGACG |
| 6hLPCAT3 Reverse(5’- 3’) | GTGGTAGAGCTGGTTTCCAAAG |
| 7hPTGS2 Forward(5’- 3’) | TCCCTTGGGTGTCAAAGGTAAA |
| 7hPTGS2 Reverse(5’- 3’) | TGGCCCTCGCTTATGATCTG |
| 8hPLCXD2 Forward(5’- 3’) | TGGCAATCCCAGGCTCACA |
| 8hPLCXD2 Reverse(5’- 3’) | ACGCAGGTCAAAGTAGCGG |
| 9hDHRS9 Forward(5’- 3’) | CATGAACTAGCAAGAGGACACCA |
| 9hDHRS9 Reverse(5’- 3’) | CAAAGCCCGAGTCACATCCA |
| LPCAT3 Promoter Primer 1 Forward | TACAAACCTTCAGACCCGCC |
| LPCAT3 Promoter Primer 1 Reverse | GCTATCGCTTCACCCCCAAT |
| LPCAT3 Promoter Primer 2 Forward | GGTGACTCCTGCCGAGAATG |
| LPCAT3 Promoter Primer 2 Reverse | TCCTGGGAGCATTTCGAACG |
| LPCAT3 Promoter Primer 3 Forward | TTACTGGGGCATGTAACGGG |
| LPCAT3 Promoter Primer 3 Reverse | GGCGACGAGTCTAACAACCT |
| LPCAT3 Promoter Primer 4 Forward | TCCCCCTAGAGGCGATGAAA |
| LPCAT3 Promoter Primer 4 Reverse | AAAGCAGGGAGGGGAAACAG |
| LPCAT3 Promoter Primer 5 Forward | CTTGCTTTGGCTGCGCAATA |
| LPCAT3 Promoter Primer 5 Reverse | TGCCCAAAATGGGAGTAGGG |
| LPCAT3 Promoter Primer 6 Forward | AGCTTCCTGAGTAGCTGGGA |
| LPCAT3 Promoter Primer 6 Reverse | CCTTATATTGGCCGGGCACA |
| LPCAT3 Promoter Primer 7 Forward | CCGTCTCAGCCTCCCAAATT |
| LPCAT3 Promoter Primer 7 Reverse | CTTGAGCCTGGGAGGTTGAG |

**Supplemental Table S3:** **The Relative siRNA in this study**

| **Gene name** | **Relative sequences (5’ - 3’)** |
| --- | --- |
| si-NPAS2-1 | TGACGAACCTTTAGAGGAA |
| si-NPAS2-2 | CAACCTCTTTGCACAGTGA |
| si-LPCAT3-1-s | GGGCUUCAAUGGCUUUGAA/dT/dT |
| si-LPCAT3-1-a | UUCAAAGCCAUUGAAGCCC/dT/dT |
| si-LPCAT3-2-s | AGGGAAAGAUCAGAAUUCC/dT/dT |
| si-LPCAT3-2-a | GGAAUUCUGAUCUUUCCCU/dT/dT |

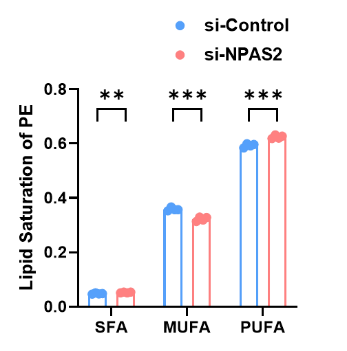

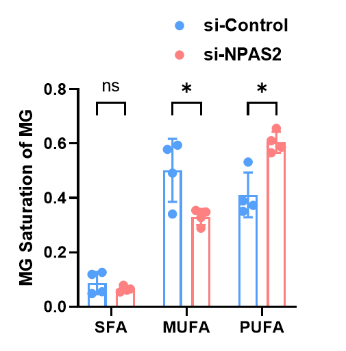

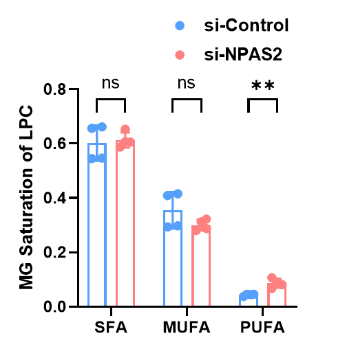

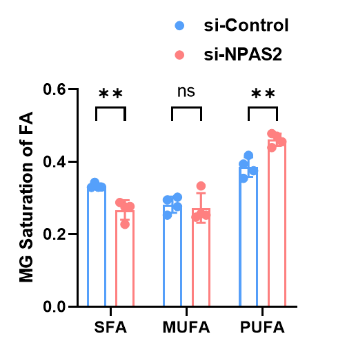

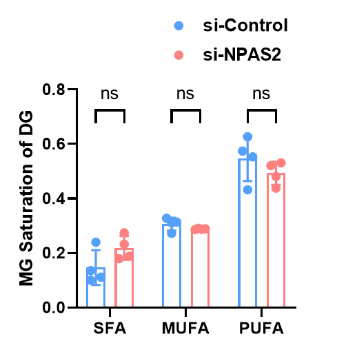

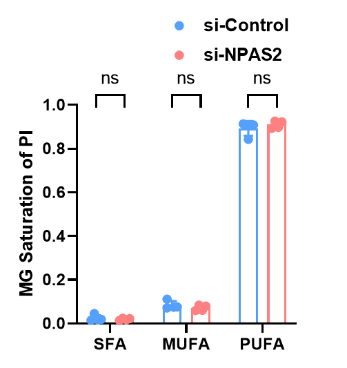

**Supplemental Figure S1: The effect of NPAS2 depletion on fatty acid saturation profiles.**
